# Astrocyte Ca^2+^ activity regulates node of Ranvier length in the white matter

**DOI:** 10.1101/2025.01.17.633566

**Authors:** Jonathan Lezmy, Tania Quintela-López, Nicholas Woermann, Shuying Mao, Caleb Thompson, David Attwell

## Abstract

Nodes of Ranvier generate action potentials along myelinated axons, but it is unclear whether they can modulate neural circuit function. Computer modelling previously predicted that adjusting node length can control axonal conduction speed^1^, but it is unknown whether mechanisms controlling nodal structure operate in the healthy brain. Here, in brain slices and in intact white matter tracts of live mice, we found that nodes elongate and shorten, while an overall stable mean node length is maintained. Changes in node length were only detected in nodes that were flanked by compact myelin sheaths and, in brains of juvenile mice, neuronal activity caused the nodes to elongate. This occurred via astrocyte Ca^2+^-mediated adenosine generation, targeting myelin sheath A_2b_ receptors which raised oligodendrocyte cyclic AMP levels. This activated NKCC1 cotransporters present in the myelin of the paranodes flanking the nodes, the expression of which was higher in the brains of juvenile mice compared to adults. NKCC1 activation elevated the membrane conductance of the myelin sheaths and reduced the length of the paranodal ends of the internodes, which was associated with node elongation that is predicted to slow action potential propagation. Thus, nodal dynamics may continuously tune information flow along myelinated axons and drive activity-dependent white matter plasticity.

## Main

The human white matter makes up about half of the human brain. It is composed of bundles of axons wrapped by oligodendrocyte-generated myelin, which reduces the axon capacitance and thus ensures rapid propagation of information between grey matter areas. In the grey matter, synapses act as units for information processing, a feature conferred by the high electrical and morphological adaptability of dendritic spines^2^. However, whether information flow is regulated in the white matter is unclear, as synapses are largely absent and information flowing along axons is essentially isolated from the extracellular space by the myelin of the internodes, except for a few unmyelinated sites: notably the nodes of Ranvier^3^. Nodes of Ranvier share characteristics of regulatory units similar to synapses because: (i) they possess the machinery to alter information encoded as action potentials^4^; (ii) the electrical properties of their membranes can be modulated in response to network activity^5^; and (iii) this modulation is mediated by cell-cell interactions, as essentially all the nodes contact astrocytes^5,6^. Just as dendritic spines can swell or shrink (or alter receptor number) to modulate synaptic integration^7,8^, nodal morphological changes could potentially control axonal excitability and conduction speed^1^. However, it is unclear whether such modulation occurs in healthy brains and what the underlying mechanisms are.

### Nodes of Ranvier are motile in healthy brains

Imaging nodes flanked by Caspr-labelled paranodes *in vivo* through a cranial window every 5 minutes for up to an hour in white matter tracts of intact Caspr-GFP mice brains (∼25 weeks old; Fig. 1a-c) revealed that 51% of them were motile (n=32/63 nodes in total; see Methods), either elongating or shortening (56% and 44% of the motile nodes, respectively; Fig. 1d-f, Supplementary Video 1 and 2). In live acute brain slices from mice of the same age (∼25 weeks old; Fig. 1e-f, Extended Data Fig. 1), similar fractions of motile nodes (45%, n=27/60; not significantly different from nodes *in vivo*: P=0.23) and of elongating and shortening nodes (63% and 37% of the motile nodes, respectively; not significantly different from nodes *in vivo*: P=0.16) were observed. Mean node length (1.68±0.05 µm and 1.72±0.08 µm, respectively; P=0.62) and node density (389,407±23,379 nodes/mm^3^ and 533,075±73,330 nodes/mm^3^, respectively; P=0.14) were not significantly different between cranial window and slice preparations (Extended Data Fig. 2). In both preparations, the bidirectional changes that we report did not alter the total mean node length over one hour (Fig. 1d, Extended Data Fig. 1d; mean node length *in vivo* in the first 15 min [1.676±0.053 µm in 63 nodes] versus last 15 min [1.681±0.054 µm in 63 nodes]: P=0.88; node length *ex vivo* in the first 15 min [1.723±0.0791 µm in 60 nodes] versus last 15 min [1.807±0.0897 µm in 60 nodes]: P=0.064). The mean node length also did not change across different ages from 2-week-old to 25-week-old mice (Fig. 2a-b; P=0.87 comparing nodes at 2, 6, 13 and 25 weeks old, one-way ANOVA).

**Fig. 1:**
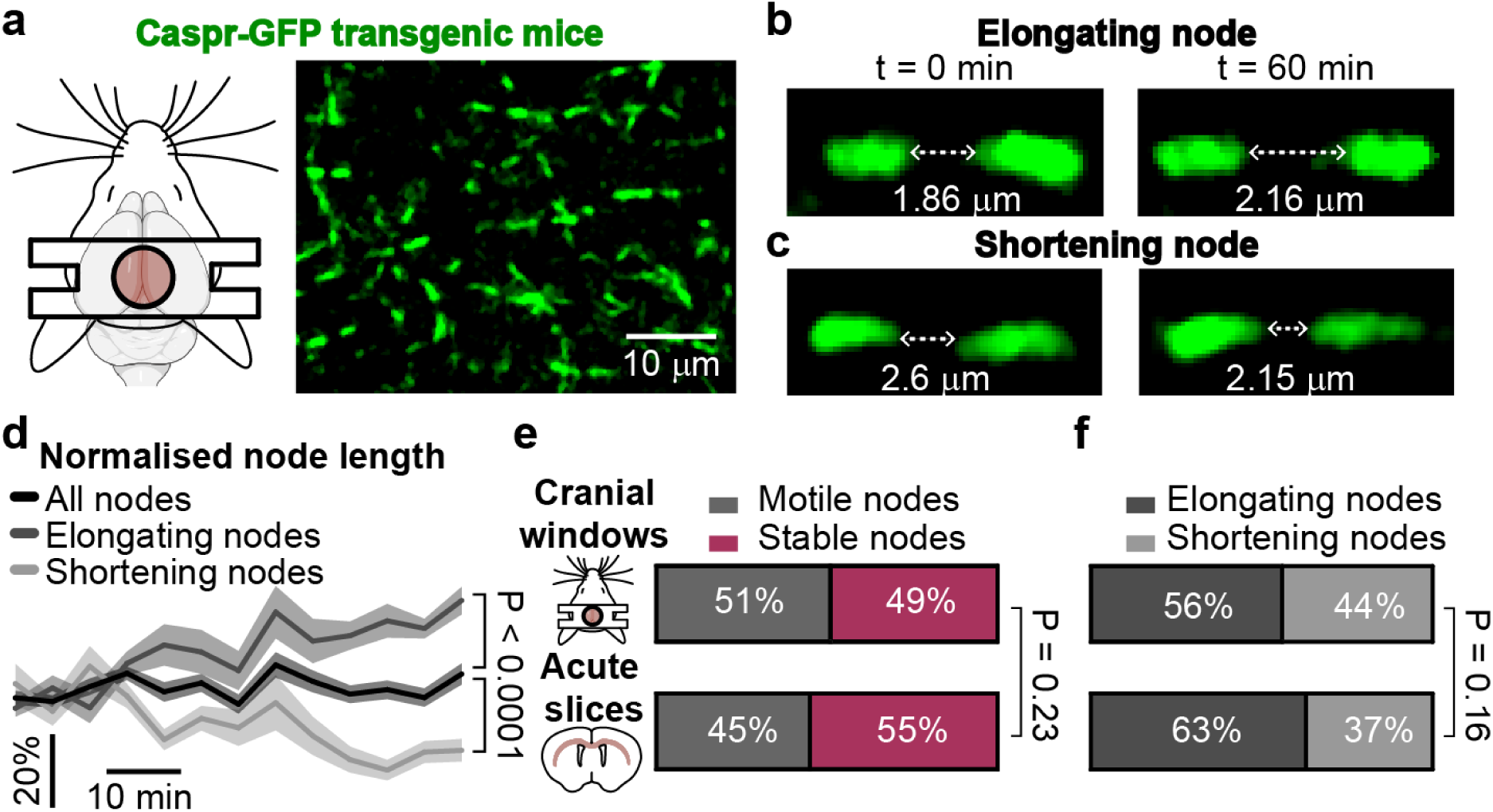
Nodes of Ranvier are motile in the healthy white matter. **a** Nodes in the white matter projections of the cingulum imaged through a cranial window in a Caspr-GFP mouse (with paranodes in green flanking nodes). **b-c** A fraction of nodes could be seen elongating (>10% longer over 1 hour, **b**) and others shortening (>10% shorter, **c**). **d** Change in node length over one hour for the elongating and shortening nodes, and for all the nodes together (*n*=18, 14 and 63, respectively). Average node length values in last 15 minutes are significantly different (P<0.0001, one-way ANOVA). **e** Percentage of motile and stable nodes identified in cingulum of mice *in vivo* through cranial windows (top) and *ex vivo* in the corpus callosum of slice preparations (bottom, not significantly different, P=0.23). **f** Percentage of elongating and shortening nodes identified in mice *in vivo* through cranial windows and *ex vivo* in slice preparations (not significantly different, P=0.16).

**Fig. 2:**
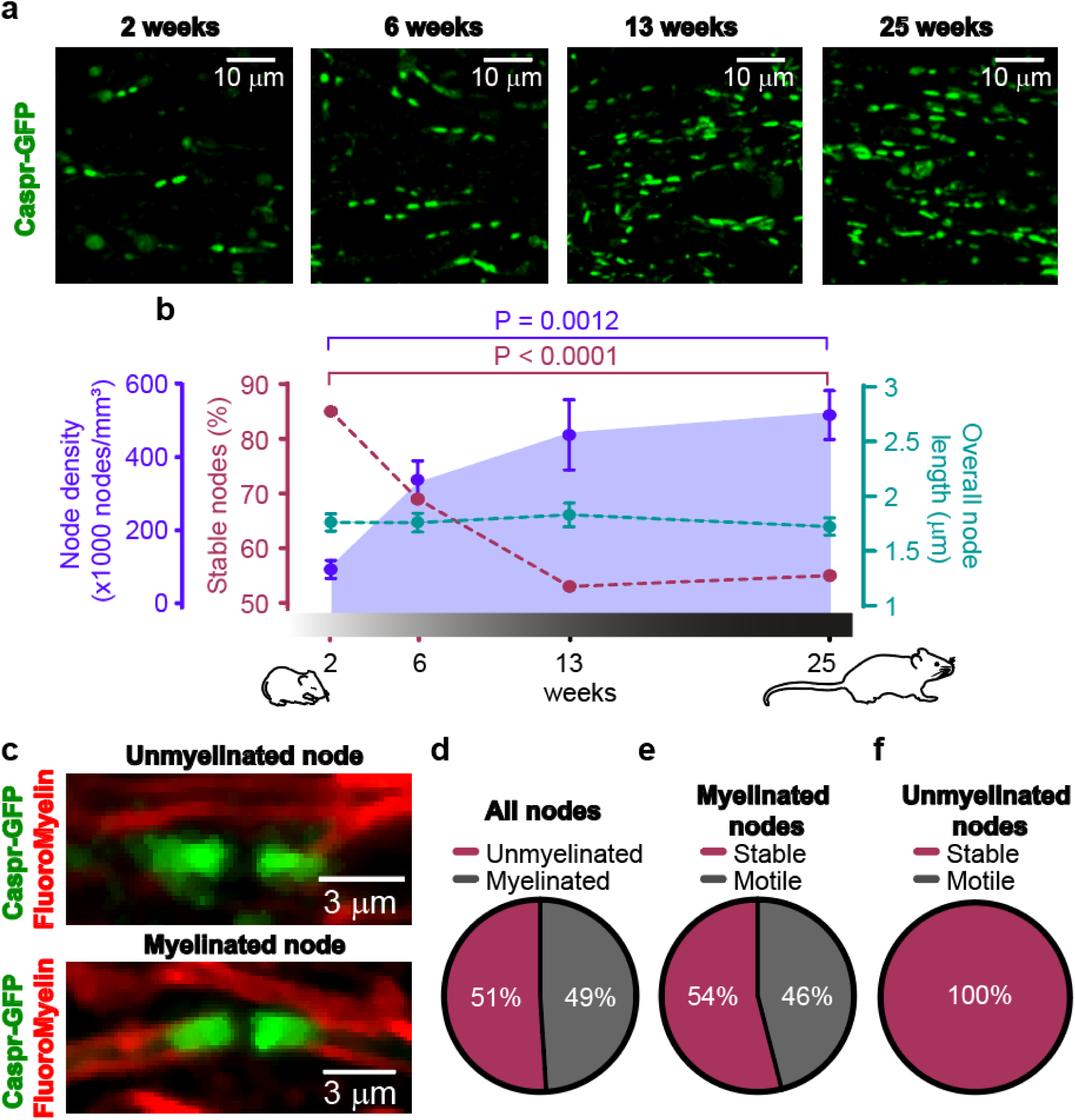
Formation of compact myelin sheaths may confer motility to nodes of Ranvier. **a** Nodes in the corpus callosum of brain slices from Caspr-GFP mice at 2 weeks old (3 mice), at 6 weeks old (3 mice), and at two timepoints in adulthood (13 and 25 weeks old, 3 mice each), but before ageing. **b** As myelination of the white matter advances, node density increases steeply, then stabilises in adulthood (2 weeks: 127,351±24,603 nodes/mm^3^, *n*=7 brain slices; 6 weeks: 368,739±51,093 nodes/mm^3^, *n*=8; 13 weeks: 489,576±94,811 nodes/mm^3^, *n*=7; 25 weeks: 533,075±65,954 nodes/mm^3^, *n*=6). Over a similar time frame, the fraction of motile nodes (100% minus the red curve data) increases steeply then stabilises (2 weeks: 85% of nodes are stable, *n*=65; 6 weeks: 69%, *n*=84; 13 weeks: 53%, *n*=51; 25 weeks: 55%, *n*=60). Mean node length remained stable across all ages (2 weeks: 1.759±0.078 µm, *n*=65; 6 weeks: 1.757±0.083 µm, *n*=84; 13 weeks: 1.829±0.083 µm, *n*=51; 25 weeks: 1.721±0.079 µm, *n*=60). **c-f** In 2 week-old mice (at which age single axons can be traced in the corpus callosum), 49% of nodes are associated with compact myelin labelled by Fluoromyelin Red (**c**, **d**). Of the nodes flanked by compact myelin, 46% were motile (**e**), and, of the nodes (defined as a region flanked by a pair of paranodes labelled with Caspr-GFP) that are not flanked by compact myelin, none were motile (**f**).

### Myelination is associated with nodes gaining motility

Having unveiled spontaneous structural motility as an attribute of nodes of Ranvier, we examined the underlying cellular mechanisms using live acute brain slices. Node density in the corpus callosum (Fig 2a-b) increased rapidly from 2 to 13 weeks old (by 362,225 nodes/mm^3^, P=0.0026), then stabilised in adulthood from 13 to 25 weeks old (increasing only by 43,500 nodes/mm^3^, P=0.97 compared to at 13 weeks), similar to the rate of myelination of callosal tracts (assessed as the increase in the fraction of myelinated axons at different ages)^9^. In parallel, the fraction of stable length nodes (Fig. 2b) decreased rapidly from 85% at 2 weeks old to 53% at 13 weeks old (P<0.0001), then remained steady in adult mice (55% at 25 weeks old). To assess the temporal changes of node motility and myelination, we imaged Caspr-GFP (to define nodes) together with FluoroMyelin Red (Fig. 2c), a marker for compact myelin^10^, in 2-week-old mice (at which age single myelinated axons in the white matter can be traced). Around half of the nodes (51%) were flanked by both Caspr-GFP-labelled paranodes and FluoroMyelin-labelled compact myelin sheaths, while the other half were flanked by Caspr-GFP only (Fig. 2d). Myelination appears to be associated with motility of nodes because, of the nodes flanked by myelin, 46% were motile (Fig. 2e) with 65% of this 46% increasing in length and 35% decreasing in length. In contrast, none of the nodes not flanked by myelin were motile (Fig. 2f).

### Node elongation depends on NKCC1 activity

Myelin remodelling is triggered by changes in brain activity, which occurs across weeks^11–13^, and may consequently induce changes in node structure, as previously reported^14,15^. However, it is unknown whether nodes of Ranvier, similar to synapses^2,16^, can adapt their shape within minutes in response to changes in neuronal activity. Enhancing neuronal activity by increasing [K^+^]_o_ from 3 to 15 mM for 45 minutes did not significantly affect node dynamics in adult mice (Fig. 3a, Extended Data Fig. 3a), but increased the fraction of elongating nodes in the white matter of young mice (Fig. 3b; from 62% to 78%, P=0.004). This effect could be prevented by inhibiting neuronal activity with 1 µM TTX or blocking the Na^+^-K^+^-Cl^-^ co-transporter NKCC1 with 20 µM bumetanide (Fig. 3b). In both young and adult mice, none of the conditions tested significantly changed the overall fraction of motile versus stable nodes (Extended Data Fig. 3a-b), possibly because roughly half of the nodes are not flanked by compact myelin and lack the ability to undergo structural changes (Fig. 2c-f). Thus, nodal plasticity dependent on neuronal activity and NKCC1 was detected in the myelinated nodes in the juvenile white matter.

**Fig. 3:**
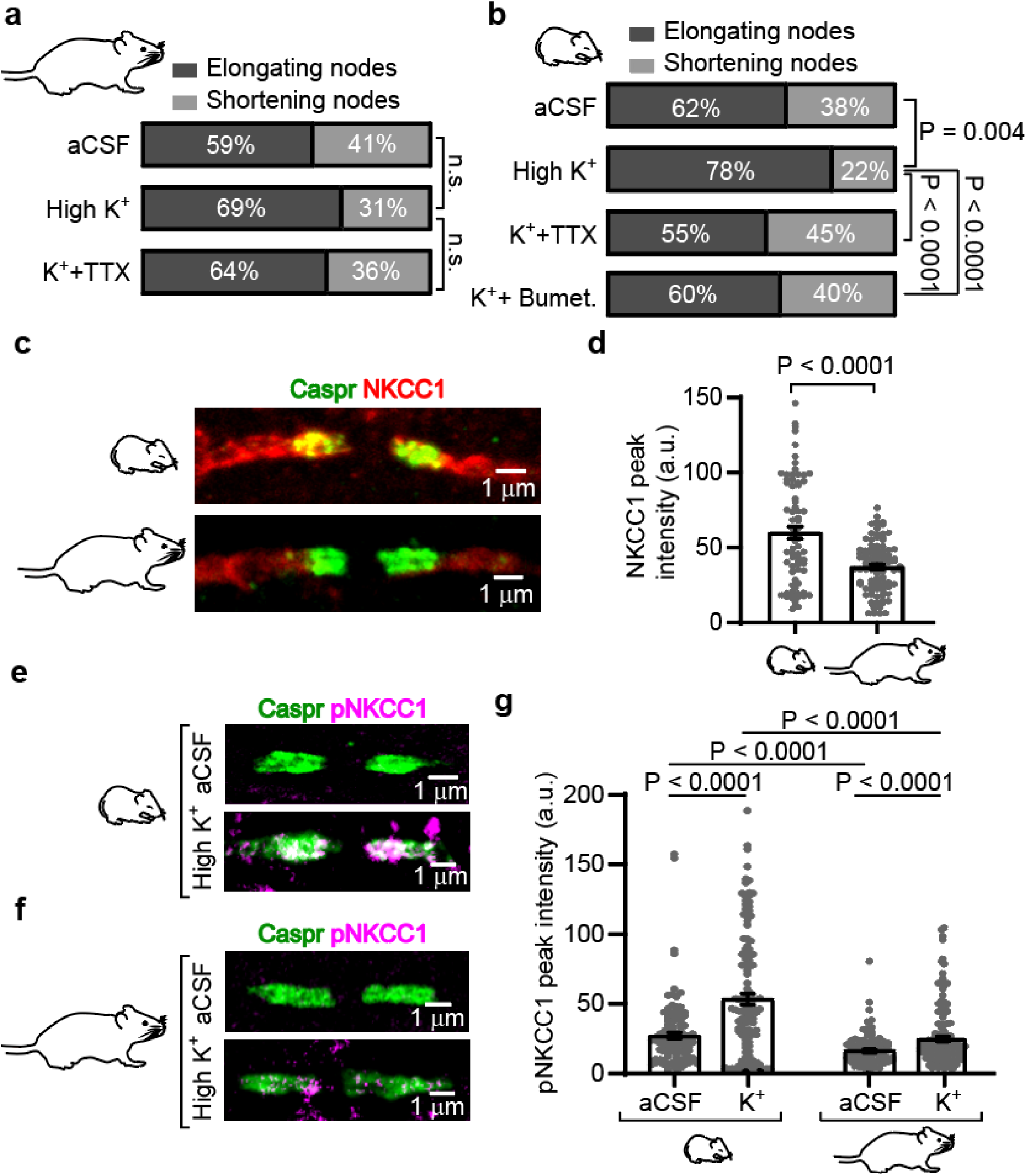
Neuronal stimulation activates NKCC1 at the paranodes and promotes the elongation of nodes of Ranvier. **a** In adult mice, the fraction of elongating and shortening nodes did not change significantly if we increased [K^+^]_o_ levels to promote neuronal activity (aCSF versus K^+^: P=0.13; K^+^ versus K^+^+TTX: P=0.56; n=59, 92 and 75 nodes in total for aCSF, K^+^ and K^+^ + TTX; n.s: non-significant). **b** In young mice, high [K^+^]_o_ increased the fraction of elongating nodes by 26%. Including TTX or bumetanide in a solution with high [K^+^]_o_ reduced by 29% and 23% the fraction of elongating nodes, respectively (n=65, 61, 55 and 65 nodes in total for aCSF, K^+^, K^+^ + TTX and bumetanide). **c** NKCC1 is present in the internode, including in the paranodal regions of young and adult mice, but its expression in adult mice appears lower (images were taken under identical illumination conditions and photomultiplier gain). **d** NKCC1 peak intensity in 75 young and 103 mature paranodal regions (young: 60.16±4.04 arbitrary units [a.u.] versus adult: 37.13±1.63 a.u; P<0.0001, unpaired *t*-test). **e** In young mice, phosphorylated NKCC1 (pNKCC1) detection (magenta, overlay with green paranodes gives white labelling) was low in paranodal regions of slices incubated in aCSF, but high when incubated in high [K^+^]_o_. **f** As in (**e**), but in adult mice; pNKCC1 levels appeared lower in adult mice treated with high [K^+^]_o_ compared to young mice. **g** Peak intensity of pNKCC1 labelling in 119 paranodal regions of young mice incubated in aCSF and 127 paranodal regions of young mice incubated in high [K^+^]_o_ (aCSF: 27.87±2.13 a.u. versus high K^+^: 54.66±3.99 a.u.; P<0.0001, two-way ANOVA), and pNKCC1 peak intensity in 90 paranodal regions of adult mice incubated in aCSF and 133 paranodal regions of adult mice incubated in high [K^+^]_o_ (aCSF: 17.37±1.22 a.u. versus high K^+^: 25.62±1.89 a.u.; all p-values are below 0.0001, two-way ANOVA). a.u.: arbitrary units.

### Neuronal activity phosphorylates paranodal NKCC1

NKCC1 transporters expressed in myelinated axons, either on the axonal or myelin membranes^17^, control myelin structure^18^ and axonal conduction^19^. NKCC1 activity regulates cell volume^20^ and intracellular [Cl^-^]_i_^21^, and its postsynaptic presence at GABAergic synapses of young brains promotes depolarising synaptic currents^21,22^. In cortical layer VI myelinated axons projecting towards the corpus callosum and in callosal axons we found, using immunohistochemistry, that NKCC1 colocalised with paranodal Caspr and internodal MBP-labelled myelin (Fig. 3c, Extended Data Fig. 4a) but was not found at nodes, suggesting that NKCC1 is preferentially expressed on the myelinating processes of oligodendrocytes (or on the underlying axonal membrane). NKCC1 was detected in 91% of paranodes in young mice (2-weeks-old) and 96% of paranodes in adult mice (25-weeks-old) (Extended Data Fig. 4b). However, NKCC1 expression, defined by its fluorescence intensity, was 1.62 times higher in the paranodes of young brains compared to adult brains (Fig. 3c-d, Extended data Fig. 4c). Compared to the internodes, NKCC1 was enriched in the paranodes of juvenile mice, where its fluorescence intensity was 1.14 times higher (Extended data Fig. 4d). The phosphorylated form of NKCC1 (pNKCC1) has a 5-10 fold enhanced co-transport activity compared to the unphosphorylated form^23^. Under physiological conditions, pNKCC1 was detected in only 11% and 13% of paranodes in young and adult mice (a detectable pNKCC1 signal was defined as being >5 times higher than background intensity; see Methods), respectively (Fig. 3e-f, Extended Data Fig. 5a). In slices incubated with high [K^+^]_o_ for 45 minutes pNKCC1 occurrence increased, being seen in 49% of paranodes in both young and adult brains (Fig 3e-f, Extended Data Fig. 5a). Paranodal pNKCC1 expression was 1.96 times and 1.47 times higher in young and adult mice following exposure to high [K^+^]_o_, but overall pNKCC1 expression (regardless of the tested conditions) was 1.87 times higher in paranodes of young mice compared to adults (Fig. 3g). pNKCC1 clustered at the paranodes of juvenile mice, where its fluorescence intensity was 1.83 times higher than in internodes (Extended Data Fig. 5b-c). Together with the effect of bumetanide reported above, these data suggest that neuronal activity promotes node elongation in the white matter via upregulation of paranodal NKCC1 activity.

**Fig. 4:**
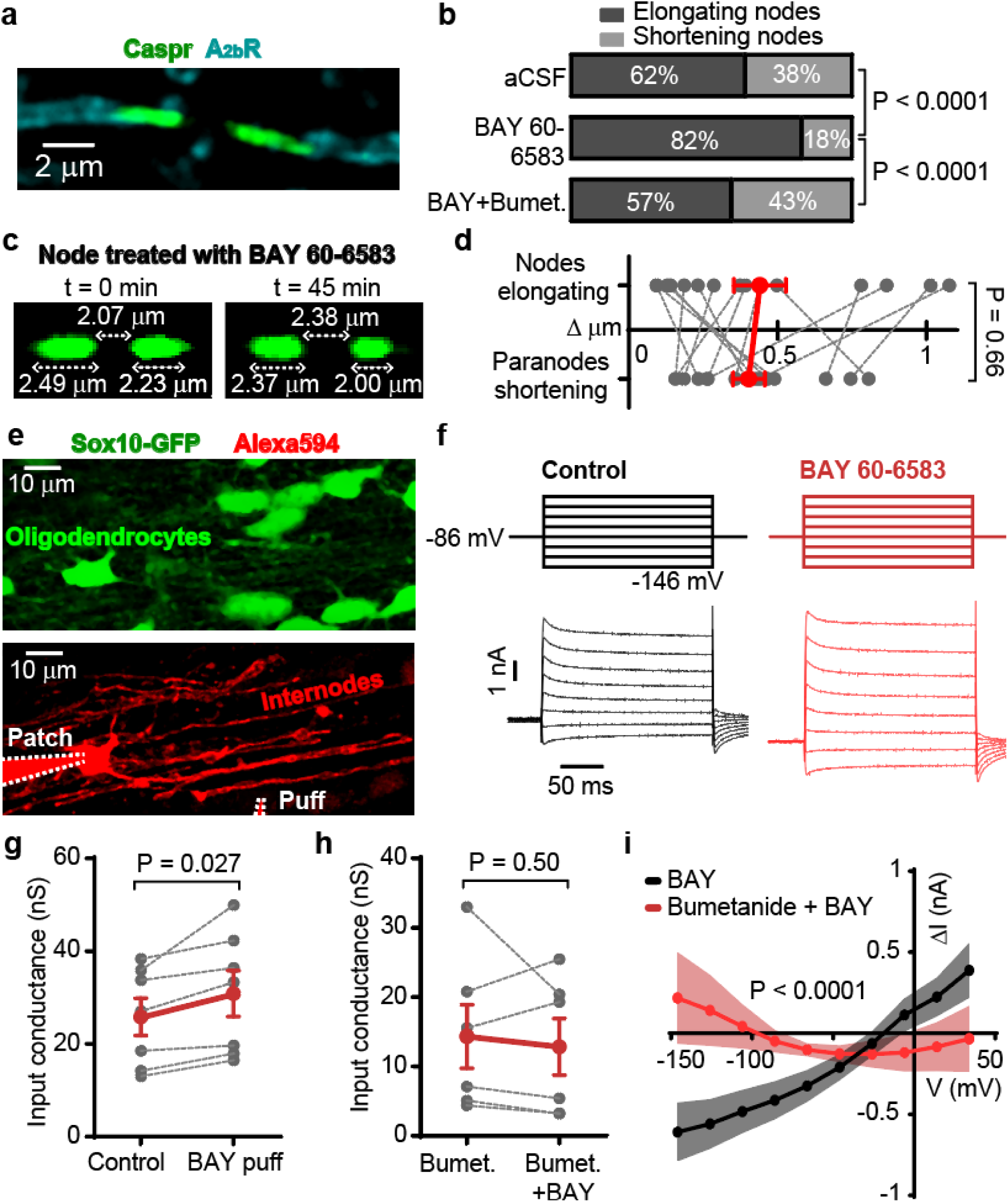
Adenosine A_2b_Rs and NKCC1 regulate node length and internodal membrane currents. **a** A_2b_Rs are present in the myelin sheaths flanking the nodes. **b** Superfusion of the A_2b_R agonist BAY 60-6583 increased the fraction of elongating nodes by 32% in young mice, similar to the effect of high [K^+^]_o_. Blocking NKCC1 with bumetanide prevented the A_2b_R-mediated nodal elongation (n=65, 87 and 50 nodes in total for aCSF, BAY 60-6583 and BAY 60-6583+Bumetanide). **c** Flanking paranodes shorten as the node elongates in a slice treated with BAY 60-6583 for 45 minutes. **d** As BAY 60-6583-treated nodes elongate, adjacent paranodes shorten to a similar extent (n=15, paired *t*-test). Each dashed line connects an observed elongation in node length to the corresponding shortening of the adjacent paranodes. Δ µm represents the absolute change in node and paranode length in slices treated with BAY 60-6583 for 45 minutes. **e** Callosal oligodendrocyte, detected in green using Sox10-GFP mice, was patch-clamped and filled in red with Alexa Fluor 594, exposing the myelinated internodes (with each internode ending at paranodes on both sides) emerging from the cell. Lower pipette was used to puff BAY 60-6583 onto myelin sheaths (see Supplementary video 3). Pipettes are delineated with dashed white lines. **f** Specimen currents elicited by +20 mV voltage steps starting from −146 mV before and after puffing BAY 60-6583, allowing the generation and analysis of I-V relations (see Methods). **g** Input conductance in aCSF before and after puffing BAY (*n*=7; P=0.027, two-way ANOVA over panels g and h). **h** As in (**g**), but while including bumetanide within the aCSF (*n*=6; P=0.50, two-way ANOVA). **i** Variations in membrane current evoked by A_2b_R activation (black curve) are prevented by NKCC1 blockade with bumetanide (red curve; P<0.0001 when comparing the slopes). Plots of subtracted currents were generated from I-V curves shown in Extended Data Fig. 7a, c.

### Myelin A_2b_ receptors regulate node motility

To explore the mechanism underlying node structural plasticity in the white matter, we focused on studying nodes in mice aged postnatal days 12 to 24 (P12-P24) that express high levels of NKCC1 in the flanking myelin sheaths. NKCC1 is phosphorylated by the cyclic AMP (cAMP)-activated protein kinase A, the latter being a target of G protein-coupled receptors coupled to G_s_^23,24^. We previously found that the G_s_-coupled adenosine A_2a_ receptors expressed at the axonal nodes, but not at the paranodes or internodes^5^, regulate axon function by modulating the electrical properties of nodal membranes^5^. Astrocyte Ca^2+^ activity mediates this effect, as it releases ATP which is converted into adenosine in the extracellular space^5^. Here, we found that another G_s_-coupled adenosine receptor, the A_2b_ receptor (A_2b_R) (Fig. 4a), is expressed in the myelin, colocalising with MBP and flanking 93% of nodes of Ranvier (Extended Data Fig. 6a-b), similar to NKCC1 (Fig. 3c-d). Astrocytes, which could also be a source of ATP/adenosine to activate these receptors, run close to the myelinated axons (Extended Data Fig. 6c). A_2b_R expression in the white matter may thus confer NKCC1 with the ability to activate in response to changes in neuronal activity, thereby mediating (or allowing) alterations in node length. Indeed, applying an A_2b_R agonist (BAY 60-6583, 500 nM) promoted node elongation (P<0.0001), mimicking the effect induced by neuronal activity, and this was abolished by bumetanide (Fig 4b; P<0.0001). In slices treated with BAY 60-6583, 85% of elongating nodes were associated with a corresponding shortening of the two adjacent paranodes (Fig. 4c-d, P=0.66; nodes elongate by 0.40±0.05 µm, the two paranodes shorten by a total of 0.44±0.09 µm), which maintained a stable overall length of the whole nodal region (a nodes with its two flanking paranodes). This data suggests that variations in node length do not require a change in the length of the compact myelin and underlying axonal membrane between the paranodes.

### A_2b_R and NKCC1 regulate myelin conductance

In the confined intracellular volume of the paranodes, NKCC1 activity will substantially increase ion concentration and osmotic water flux, potentially leading to changes in membrane currents. To assess the electrical effect of A_2b_R and NKCC1 on the myelin membrane, we patch-clamped myelinating oligodendrocytes, targeting cells in cortical layer VI or the corpus callosum of Sox10-GFP transgenic mice (Fig. 4e). When puff-applying 500 nM BAY 60-6583 onto myelin sheaths (Fig. 4f-g, Extended Data Fig. 7a-b, Supplementary Video 3), the input conductance of oligodendrocytes (*g*_in_) increased from 25.8±4.0 nS to 30.9±5.0 nS (P=0.027) and the membrane resting potential (*E*_m_) depolarised from −76.6±4.3 mV to −65.4±4.5 mV (P=0.029). Puffing artificial cerebrospinal fluid (aCSF; Extended Data Fig. 7f) had no significant effect on the current-voltage (I-V) relationship (Δ*g*_in_=-1.3±1.7 mS and Δ*E*_m=_+2.8±2.2 mV; I-V relation slope not significantly different: P=0.87). Adding 50 µM intracellular cAMP to the oligodendrocytes via the patch-pipette (Extended Data Fig. 8), which spread to the myelin sheaths, produced an effect on the membrane current similar to that of A_2b_R as it increased *g*_in_ by +6.2 nS and depolarised *E*_m_ by +5.2 mV (p<0.0001, Extended Data Fig. 8d; data were derived from the linear regression fitted to the I-V relations) and, in these oligodendrocytes, BAY 60-6583 did not have a significant further effect on the membrane current (Extended Data Fig. 8a-c; Δ*g*_in_=-1.5±1.3 nS and Δ*E*_m_=+1.9±1.1 mV; P=0.33 and P=0.55, two-way ANOVA comparing with and without puff-application of BAY 60-6583 when cAMP is included in the patch pipette). Superfusion of 20 µM bumetanide alone (Extended Data Fig. 7e) reduced the input conductance by 11.8 nS (P<0.0001, although *E*_m_ only slightly hyperpolarised by −0.8 mV), which may reflect basal levels of NKCC1 endogenous activity in myelin sheaths (as shown also by the detection of pNKCC1 in some paranodes in control conditions; Fig. 3g, Extended Data Fig. 5a-b). The presence of superfused bumetanide prevented the generation of the A_2b_R-triggered membrane current when BAY 60-6583 was puffed onto myelin sheaths (Δ*g*_in_=-1.5±2.5 mS and Δ*E*_m_=+0.9±5.2 mV; P=0.50 and P=0.86, two-way ANOVA comparing with and without puff-application of BAY 60-6583 in aCSF containing bumetanide; Fig. 4h-i, Extended Data Fig. 7c-d).

### A_2b_R and NKCC1 activity shorten paranode length

To understand the association between structural and electrical variations in myelin properties, patch-clamped oligodendrocytes were dye-filled with Alexa Fluor 594, revealing the myelinated internodes emerging from a single cell, with each internode ending at paranodes on both sides (Fig. 4e, 5a). Within 10 minutes of applying BAY 60-6583 (Fig. 5b), the length of the internodes targeted by the puff (see Methods and Supplementary Video 3) decreased from its mean value of 26.5 μm by 5.99±1.82% (P=0.003) but the length of the internodes located away from the puffed area (Extended Data Fig. 9, see Methods) was not affected (−0.61%±1.93%; P=0.76, *n*=6). This corresponds to each paranode shortening by 0.79±0.24 µm (half the total shortening because there are two paranodes at the ends of the internode). This would result in a node elongation from 1.76 µm (the mean node length reported in Fig. 2b) to 3.34 µm. When NKCC1 was blocked with superfused bumetanide (Fig. 5c), no change in internode length was observed after puffing BAY 60-6583 onto the myelin sheaths (−1.80%±1.32%; P=0.015 from a 2-way ANOVA when compared with puffing BAY 60-6583 without bumetanide in the aCSF). Thus, node elongation mediated by A_2b_R and NKCC1 activity (Fig. 3b, 4b) was observed with the shortening of the Caspr-GFP labelled paranodes (Fig. 4c-d) and at the paranodal ends of dye-filled myelinated internodes (Fig. 5b). A longitudinal shortening of the internodes at their ends where the paranodes are located (since pNKCC1 is clustered in these regions; Fig. 3e-g), might be expected to increase the length of the nodes flanked by these paranodes. Paranodal loops, with their high cytosolic content compared to the compacted myelin lamellae along the remaining length of the internode, may also be more likely than the main internode to accommodate intracellular signalling cascades involving cAMP rises and NKCC1 phosphorylation, and thus be more susceptible to length changes.

**Fig. 5:**
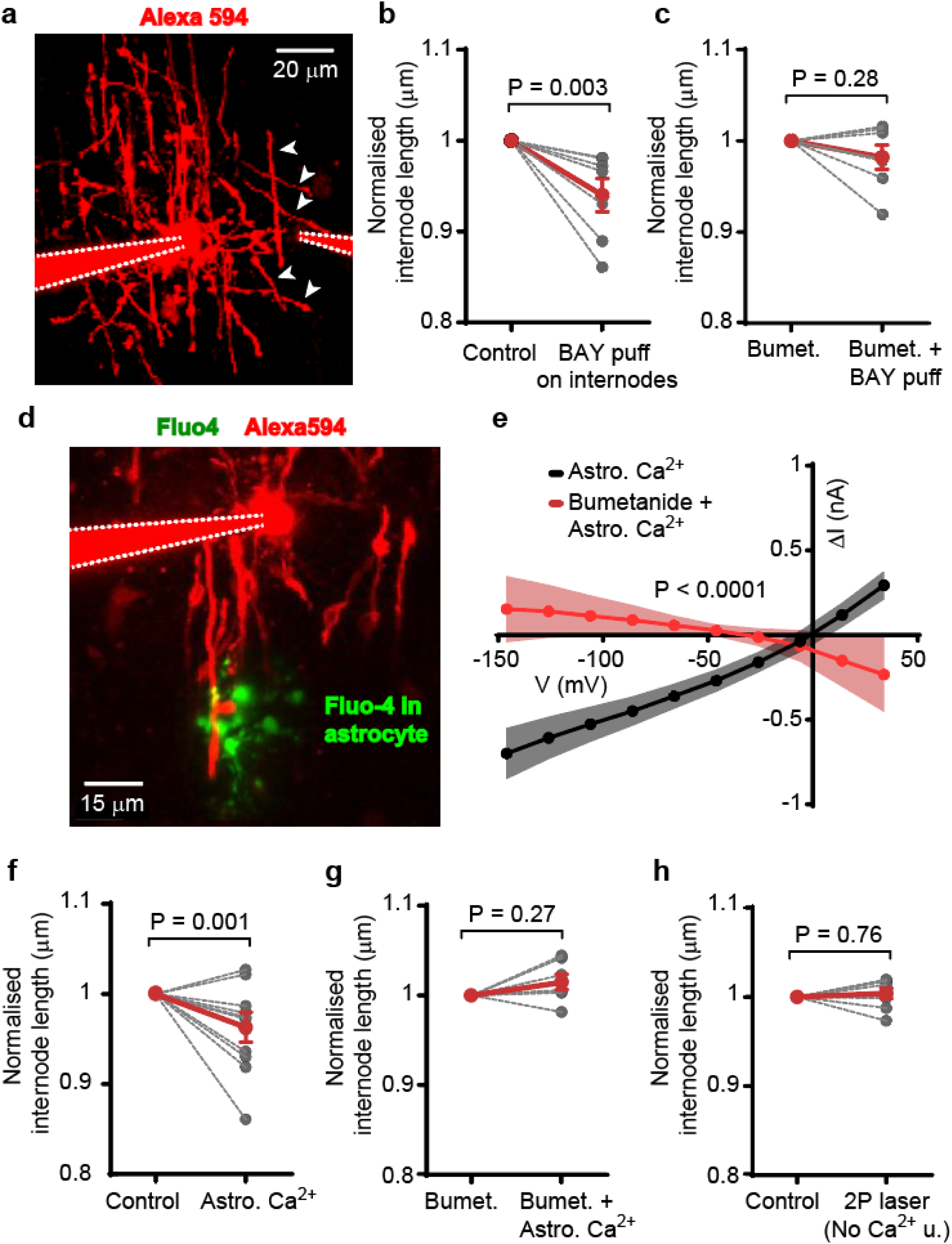
Astrocyte Ca^2+^ activity near myelin sheaths regulates internode length and oligodendrocyte membrane currents. **a** Layer VI oligodendrocyte patched-clamped and dye-filled with Alexa 594 in the left pipette. The right pipette was used to puff the A_2b_R agonist BAY 60-6583 onto the labelled internodes. Pipettes are delineated with dashed white lines and the white arrowheads show internodes targeted by the local application of the drug (see Methods and Supplementary video 3). **b** Normalised changes in internode length (with each internode ending at paranodes on both sides), when BAY 60-6583 is puff-applied onto internodes (*n*=7, two-way ANOVA over panel b and c). **c** As in (**b**), with bumetanide superfused in aCSF prior to BAY 60-6583 application (*n*=7, two-way ANOVA). **d** Patch-clamped layer VI oligodendrocyte (in red, left pipette delineated with white dashed lines) and astrocyte filled with the Ca^2+^ cage NP-EGTA and the Ca^2+^ indicator Fluo-4 (green; see also Supplementary video 4) to raise and monitor [Ca^2+^]_i_ near myelin sheaths. **e** Variations in membrane current evoked by astrocyte [Ca^2+^]_i_ rises (black curve) are prevented by NKCC1 blockade with bumetanide (red curve; P<0.0001 when comparing the slopes). Plots of subtracted currents were generated from I-V curves shown in Extended Data Fig. 10a, d. **f-g** Normalised changes in internode length after astrocyte Ca^2+^ uncaging without (**f**; *n*=11, two-way ANOVA over panels f and g) or with bumetanide in the aCSF (**g**; *n*=7, two-way ANOVA). **h** Two-photon laser illumination did not evoke a change in internode length if it failed to elicit a Ca^2+^ rise (No Ca^2+^ u.; *n*=8, two-way ANOVA).

### Astrocyte Ca^2+^ triggers internodal adaptations

Astrocyte processes associate with internodes and nodes^5,6^, running parallel to myelinated axons in the white matter (Extended Data Fig. 6c). We found that the mean node-astrocyte process distance (from the centre of a node to the closest astrocyte process) in the corpus callosum was 0.57±0.04 µm (*n*=221). Because neuronal activity triggers astrocyte Ca^2+^ activity^26,27^, which promotes the release of ATP from astrocyte processes, and ATP is converted extracellularly to adenosine^5,28^, we elicited astrocyte Ca^2+^ transients near myelin sheaths. We patch-clamped astrocytes to uncage Ca^2+^ (see Methods and Supplementary Video 4), including the Ca^2+^ cage NP-EGTA and the Ca^2+^ indicator Fluo-4 in the patch-pipette. Another patch-pipette on oligodendrocytes was used to record their membrane currents and image their myelinated internodes dye-filled with Alexa Fluor 594 (red in Fig. 5d). Within 12 minutes of raising astrocyte [Ca^2+^]_i_ with 2-photon laser excitation (Fig. 5e, Extended Data Fig. 10a-c), the input conductance of oligodendrocytes increased by 5.3±1.0 nS (P<0.0001) and their resting potential depolarised by 9.7±2.5 mV (P=0.0004), but this was prevented by blocking NKCC1 with superfused bumetanide (Fig. 5e, Extended Data Fig. 10d-f). When the 2-photon laser did not trigger a rise in astrocyte [Ca^2+^]_i_, no change in oligodendrocyte membrane current was observed (Extended Data Fig. 10g-i). In parallel, the length of the myelinated internodes shortened by 3.92%±1.44% from a mean value of 25.62 μm (P=0.001; corresponding to each paranode shortening by 0.50±0.18 µm because there are two paranodes per internode) after triggering astrocyte Ca^2+^ transients (Fig. 5f), but this was not observed when NKCC1 was blocked or when the 2P laser power was too low to raise astrocyte [Ca^2+^]_i_ (P=0.27 and P=0.76 from 2-way ANOVAs, respectively; Fig. 5g, h). Astrocyte Ca^2+^ activity thus recapitulates the electrical and structural changes mediated by A_2b_R and NKCC1 expressed on paranodes.

### Paranodal adaptations slow conduction speed

We used an infinite cable model^1,5,29^ to predict the effect of myelin A_2b_Rs and NKCC1 on axonal conduction speed in the corpus callosum (see Methods). We separately tested the effects of two results obtained from our experimental data: the rise in myelin conductance and node elongation. To estimate the process conductance and the specific conductivity of internodal membrane from patch-clamping oligodendrocytes, we made several assumptions, including that current passage to the extracellular space across the membranes of an oligodendrocyte’s myelin processes occurs solely across the outer myelin membrane (because the inner membranes are too electrically isolated; see Methods). Input conductance correlated significantly with oligodendrocyte internodal process length, suggesting that the processes are electrically coupled to the soma^29^. We then derived from our patch-clamping experiments the mean process conductance (44.88 pS/µm), and its rise tiggered by A_2b_R activation (108.4 pS/µm, as defined by the slope of the linear regression in Extended Data Fig. 11a-b; P=0.021 and P=0.0013, respectively: see Methods). Based on these values and the geometrical properties of the myelin, we estimated that, when A_2b_ receptor activation occurs, then in the computer model the specific conductivity of the internodal myelin membranes has to be set to rise from 15.86 pS/μm^2^ to 16.97 pS/μm^2^ in order to represent the overall change of resistance across the whole myelin sheath. This estimated rise in myelin conductance density, if it occurs in myelin sheaths all along the axon (e.g. in conditions when adenosine is generated over an extended spatial area), is expected to only marginally slow down the conduction speed (by 0.3%; Extended Data Fig. 11c, see also Methods). We then simulated a longitudinal shortening of the paranodes, implying that the internodes shorten at their paranodal ends and the nodes elongate accordingly, as observed experimentally (Fig. 4c-d, 5b). These changes are expected to maintain an overall constant axonal length and, if they are not accompanied by the insertion of channels at the nodes, to retain the same number of voltage-gated channels at nodes (see Discussion and Methods). Increasing node length this way (rather than by keeping a constant channel density) all along the axon from the shortest to the longest observed node suggests that a decrease of conduction speed by 19.7% could occur (Extended Data Fig. 11d). In contrast, if the node channel density is kept constant, simulating the insertion or removal of membranes with channels at the nodes (which may occur during node formation, see Discussion), such an increase in node length would speed up the conduction by 21.3% (Extended Data Fig. 11d). Thus, our simulations predict that adenosine release onto myelinated axons can significantly alter the conduction speed by elongating node length, and shortening the myelinated paranodes.

## Discussion

Complex processing of neural information occurs in the grey matter but, to obtain a comprehensive understanding of brain networks, it is also important to understand the mechanisms modulating information flow along white matter projections. Our data (summarised in Extended Data Fig. 12) reveal that nodes of Ranvier are plastic structures in the white matter that can generate rapid and bidirectional variations in axonal conduction speed. We made five main findings that contribute to our understanding of the role of nodes as regulatory units: (i) they constantly vary their length in the healthy brain; (ii) the presence of myelin at paranodes is correlated with an ability to adjust node length; (iii) myelin A_2b_Rs and NKCC1 transporters regulate node length; (iv) these proteins regulate myelin sheath conductance and internode length at their paranodal ends; and (v) neuronal activity and astrocyte Ca^2+^ activity trigger the observed changes at nodes and paranodes. Together, these data indicate that node of Ranvier plasticity, via structural changes at their flanking myelinated paranodes that are triggered by associated astrocytes, provides a potential means to adjust spike propagation along myelinated axons.

Changes in conduction speed imply changes in spike arrival time at the axons’ output synapses, thereby shaping neural circuit function^30^. In this work, we identified that neuronal and astrocyte activity impact conduction speed largely by increasing node length, which increases the capacitive load of nodal membranes and thus will impede spike generation (although if more Na^+^ channels are inserted when the node elongates then the conduction speed can be increased). Our data indicate that node elongation is associated with a shortening of the paranodal ends of the internodes, conceivably exposing unmyelinated axonal membrane at the nodes without altering nodal ion channel numbers. The number of nodal Na_v_1.6 channels has previously been reported to correlate with node length^1^, possibly reflecting a relatively constant channel density across nodes when they are being formed. During their formation, nodal components, including Na_v_ channels, ankyrin-G and β4-spectrin, do not require the presence of intact myelinated paranodes to cluster at nodes^31,32^. Thus, inserting channel-containing membranes, possibly via exocytosis of vesicles, is likely to set the baseline length of the forming nodes to approximately optimise the axonal conduction speed (as seen in Extended Data Fig. 11d). While the node length observed and the myelin conductance estimated from our experimental data are set to generate almost the maximum conduction speed possible, modulation of node length by A_2b_R/NKCC1 is expected to operate in a range of values that trigger significant variations of conduction speed (Extended Data Fig. 11d). We predict that the maximum A_2b_R-evoked effects (provided all paranodes undergo the same changes simultaneously) can reach a 1.25-fold reduction of conduction speed. Together with the estimated 1.91-fold decrease of conduction speed generated by A_2a_R activation at nodes^5^, extracellular adenosine should be able to slow down the conduction speed up to 2.39-fold along a single myelinated axon. Thus, adenosine signalling represents a powerful modulator of white matter information flow.

Despite the potentially robust and widespread impact of adenosine signalling in the white matter, our data imply that influences governing spike propagation along myelinated axons are complex. Release of ATP/adenosine triggered by astrocyte Ca^2+^ activity raises oligodendrocyte input conductance by an average of 5.28 nS. If the A_2b_R/NKCC1-mediated rise in input conductance per unit length of myelinated internode is 108.4 pS/µm (as calculated from our experimental data; Extended Data Fig. 11b, see also Methods), this 5.28 nS rise corresponds to activating A_2b_Rs on internodal processes with a total length of 48.71 µm. This suggests a localised influence of an astrocyte on no more than one full length of a mouse callosal internode^33^, potentially affecting only a single node of Ranvier, which may be sufficient to fine-tune the timing of an action potential’s arrival at an output synapse by 60 µs (estimated using our computer model), i.e. within a time window that promotes neuronal plasticity^34,35^. Alternatively, if the released adenosine does not saturate myelin A_2b_Rs, the 5.28 nS rise in oligodendrocyte input conductance by astrocyte-mediated adenosine may occur over longer lengths of internodal processes (and thus possibly more intervening nodes). Since these receptors have a low affinity for adenosine^36^, their endogenous activation may occur only where there is tight physical association between myelin sheaths and astrocyte processes^5,6^. Thus, assuming a robust ATP vesicular release from astrocytes near axons during prolonged neuronal activity^5^, structural adaptations at the axons may be dictated by the topography of astrocyte processes around internodes and nodes. Astrocytic ATP/adenosine signalling is particularly prominent during periods of high energy demand, such as during wakefulness, cognitive tasks or development, which are characterised by intense neuronal activity^37,38^. These conditions may physically enhance astrocyte process association with nodes, just as they ensheathe synapses to shape neural networks^39^. Thus, the mechanisms evoked by astrocyte Ca^2+^ activity can provide precise and efficient modulation of white matter information flow along a single axon.

Transient expression of cation-chloride cotransporters and adenosine receptors at GABAergic synapses plays a crucial role in the development of neural circuits^21,40^. GABA receptors are also present in myelinating oligodendrocytes where they are coupled to the activity of NKCC1 cotransporters^19,25,41,42^, and our data demonstrate that NKCC1 and its activated phosphorylated form are found in the myelin of developing brains. We suggest that these proteins sense neuronal activity to tune spike propagation in the white matter, effectively shaping neural network function across different brain areas. Despite lower levels of NKCC1 in adult brains^21,22^ that may lessen their ability to generate changes in oligodendrocytes during brain activity, node motility was not lost in adult mice. Although we cannot rule out the existence of other mechanisms regulating node length in adult brains, NKCC1 activity, even at low levels, may still modulate node length, for instance in response to oscillatory firing^43^, causing repeated cycles of shrinkage and swelling of the myelin sheaths, as NKCC1 becomes phosphorylated when cells decrease in volume^23^. This may allow rapid and efficient modulation of the speed of information transfer, in contrast to the time and energy required for formation or removal of myelin sheaths in the dense white matter^1,11,44^. Our results thus suggest that the nodal region is a key site for altering information flow in the white matter. Many neurological disorders, such as epilepsy, schizophrenia and autism have been linked to impairment in synaptic integration, which can involve malfunction of NKCC1 and adenosine receptors^45,46^. Our work raises the question of to what extent brain malfunction in these diseases is caused by dysfunctional plasticity mechanisms operating in the white matter.

## Methods

### Animals

Transgenic mice housed on a 12 h/12 h light/dark cycle were used in all experiments. We assessed changes across different ages using mice at P12-P180 (before ageing occurs^47^). For electrophysiology experiments, we used mice at the age of P16-P24, as A_2b_R- and NKCC1-dependent node of Ranvier elongation occurred around that age. Each experiment was performed on brain slices from at least three animals and at least one of each sex. For major results, we found no differences between the two sexes: for example, in the expression of A_2b_Rs and NKCC1 in nodal regions (9.8% higher for females, P=0.15, and 3.2% higher for males, p=0.45, respectively). Thy1-Caspr-GFP mice (named Caspr-GFP in the main text), which were used to monitor node length, were generated as previously described^48^. Sox10-lox-GFP-STOP-lox-DTA mice (named Sox10-GFP in the main text), which were used to target oligodendrocytes for patch-clamping, were generated as previously described^49^. Animal procedures were carried out in accordance with the guidelines of the UK Animals (Scientific Procedures) Act 1986 and subsequent amendments (under Project Licence PP1909687).

### Acute brain slice preparation

Coronal cortical slices (300 μm thick) were prepared on a vibratome in ice-cold solution. For node length imaging, a standard slicing solution was used containing (in mM): 1.25 NaH_2_PO_4_, 2.5 KCl, 7 MgCl_2_, 25 NaHCO_3_, 25 glucose, 87 NaCl, 75 sucrose, and 0.5 CaCl_2_. The slices were incubated at 37°C in artificial cerebrospinal fluid (aCSF) for 20 min, then incubated at room temperature until use. For electrophysiology, a slicing solution adapted to achieve stable recordings contained (in mM): 93 N-methyl-D-glucamine (NMDG) chloride, 2.5 KCl, 30 NaHCO_3_, 10 MgCl_2_, 1.2 NaH_2_PO_4_, 25 glucose, 0.5 CaCl_2_, 20 HEPES, 5 sodium ascorbate, 3 sodium pyruvate and 1 kynurenic acid. The slices were incubated at 37°C in this solution for 20 min, then transferred to a similar solution with (in mM): 93 NaCl, 1 MgCl_2_ and 2 CaCl_2_ instead of the NMDG chloride, MgCl_2_ and CaCl_2,_ and incubated at room temperature until use. All experiments were performed in aCSF containing (in mM): 125 NaCl, 3 KCl, 26 NaHCO_3_, 2 MgCl_2_, 2 CaCl_2_, 1.25 NaH_2_PO_4_ and 10 glucose, heated to 37°C. All the solutions were gassed with 95% O_2_ and 5% CO_2_. In some experiments, 20 μM bumetanide (Cayman Chemical Company) was added to the aCSF or KCl concentration was increased to 15 mM instead of 3 mM (a 12mM or 24mOsm difference). To prevent effects triggered by osmolarity changes, osmolarity of control solutions with low [K^+^] was adjusted with 24 mM sucrose to match solutions with high [K^+^].

### Cranial window preparation

Caspr-GFP mice aged around P180 were anesthetised by intraperitoneal administration of ketamine (100 mg/kg) and xylazine (10 mg/kg), and anaesthesia was maintained by intramuscular injections of the same compounds. Adequate anaesthesia was ensured by confirming the absence of a withdrawal response to a paw pinch. Body temperature was maintained at 36.8°±0.3°C and eyes were protected from drying by applying polyacrylic acid eye drops (Dr. Winzer Pharma). The animal was secured in a stereotaxic frame and lidocaine/prilocaine (AstraZeneca) was applied topically prior to exposing the skull. A custom-built headplate was then attached to the skull using superglue to create a sealed well filled with HEPES-buffered aCSF (140 mM NaCl, 10 mM HEPES, 2.5 mM KCl, 1 mM NaH_2_PO_4_, 10 mM glucose, 2 mM CaCl_2_, and 1 mM MgCl_2_). A craniotomy of approximately 3 mm diameter was performed over the area where the cingulum projection axons are the closest to the brain surface^50^, along the sagittal suture and immediately rostral to the lambda point. The dura was left intact to reduce perturbation of the brain. Agarose (1%) covered the exposed brain and kept in place a glass coverslip used for imaging. During imaging, the headplate was secured under the objective on a custom-built stage.

### Live-imaging nodes of Ranvier

Through cranial windows, nodes were imaged using two-photon microscopy on a Zeiss LSM710, with a 20x water immersion objective (W Plan-Apochromat, NA 1.0). Two-photon excitation was carried out using a Newport-Spectraphysics Ti:sapphire MaiTai laser pulsing at 80 MHz. Fluorescence was evoked using a wavelength of 920 nm and the mean laser power under the objective did not exceed 35 mW. Image z-stacks were taken in 1-µm depth increments in the cingulum just below the cortical layers (up to 500 µm deep from the cortical surface). In brain slices, nodes were imaged using confocal microscopy on a Zeiss LSM780, with a 20x water immersion objective. Excitation was performed with an argon laser using a wavelength of 488 nm and laser power did not exceed 2% of its maximum (i.e. <0.6 mW under the objective). In both *in vivo* and *ex vivo* experiments, the nodes and axons were approximately aligned with the x-y plane. Image z-stacks were taken in 1 µm depth increments in the corpus callosum. Nodes of Ranvier were imaged for 0.67 seconds every 5 minutes, for 45-60 minutes.

### Quantification of node of Ranvier and paranode length

The maximum intensity of the z-stack images was projected using ImageJ (FIJI). When node density was high, we avoided overlap of Caspr signals from paranodes at different depths in the slice by projecting the maximum intensity of a subset of the z-stack, splitting the original acquisition into 5-10 z-stack images. A line was drawn along the axon, spanning both Caspr-labelled paranodes to plot the Caspr intensity profile across the node (the width of the line was adjusted to fit the width of Caspr). The size of the node was then calculated using a MATLAB script which measures the distance between the half-maximum intensity points where the end of each paranode meets the node. Paranode length was calculated using a MATLAB script which measures the distance between the half-maximum intensity points at both ends of each paranode. Nodes and whole nodal regions (a node with its 2 paranodes) were considered elongating or shortening if their length changed by >10% in 45-60 minutes. To quantify spontaneous node motility in physiological conditions, nodes were imaged every 5 minutes for 1 hour, and the mean node length of the last 15 minutes was compared to the mean node length of the first 15 minutes. To quantify the effect of different drugs or conditions on node motility, nodes were imaged every 5 minutes for 45 minutes and the node length at the end of the experiment was compared to the node length before application of the tested condition. Node density was calculated by counting nodes within a volume of interest with an area of 85×85 µm^2^ and a depth between 10 µm and 49 µm (11-50 z-stacks). Absolute node density may be underestimated as GFP may not label all the nodes in the Caspr-GFP mice (the recombination of Cre recombinase is expected to be 60-90%^51^).

### Patch-clamping of oligodendrocytes

Experiments were performed using a Zeiss LSM780 two-photon/confocal microscope, with a W Plan-Apochromat 20x objective. Myelinating oligodendrocytes in cortical layer VI or the corpus callosum were identified by their green labelling in Sox10-GFP mice. Sox10 is also expressed in oligodendrocyte precursor cells but myelinating oligodendrocytes can be distinguished as they have a distinct morphology with processes aligned with axons and they display essentially ohmic currents in response to voltage steps. Microelectrodes with resistances of 5-6 MΩ were pulled from borosilicate glass capillaries (Harvard Apparatus) and filled with an intracellular solution containing (in mM): 145 K-gluconate, 2 MgCl_2_, 0.5 H_2_-EGTA, 2 MgATP, 0.2 Na_2_GTP, and 10 HEPES, pH adjusted to 7.2 with KOH. Somata were patch-clamped in whole-cell configuration, and the signals were amplified using a Multiclamp 700B (Molecular Devices), filtered at 4 kHz and digitised at 50 kHz. The series resistance was <20 MΩ, was compensated by ≥60%, and did not vary by >10% during experiments. Recordings were acquired with Clampex and analysed with Clampfit software (Molecular Devices). All membrane potentials were corrected for a pipette liquid junction potential of −16 mV.

### Quantification of membrane currents

To build I-V curves, we patch-clamped the cells at −86 mV and then recorded currents at different voltage steps from −146 mV to +34 mV in 20 mV increments for 500 ms. We plotted the mean current in the last 100 ms of each voltage step. We retrieved two parameters from the I-V relations: the input conductance (*g_in_*) being the slope of the curve, and the resting potential (*E*_m_) being the intercept with the x-axis. If the data were acquired from the same cells (paired data), *g_in_* and *E*_m_ for each cell were retrieved from each of their I-V relations, then averaged and compared before and after a tested condition. If the data were acquired from two different sets of cells (unpaired data), average I-V relations for each set of cells were first plotted, then the mean values of *g_in_* and *E*_m_ were retrieved from the plots for the two sets of cells and were compared.

### Quantification of internode length

The myelinated internodes emerging from a single oligodendrocyte, with each internode ending at paranodes on both sides, were detected by adding 100 μM Alexa Fluor 594 into the intracellular solution of the patch-pipette targeting the oligodendrocyte soma, and the diffusion of Alexa Fluor 594 into the myelin sheaths was monitored live. A line was drawn along myelinated internodes that could be fully traced and that had clearly defined ends, and thus our selection may have favoured short internodes. Internode length ranged from 8.01 µm to 47.90 µm and averaged 25.30 µm. Intensity profiles were plotted along the internodes, the width of the line used was adjusted to fit the internode width, and the length of the internode was defined manually from the points where the intensity dropped to half of the maximum intensity.

### Local application of drugs to myelin sheaths

aCSF, alone or with BAY 60-6583 (500 nM, Tocris Bioscience), was puff-applied onto myelin sheaths dye-filled with Alexa Fluor 594. The solutions in the puff-pipette also included 100 μM Alexa 594 to monitor the spread of the puffed solution. Pipettes with a resistance of 2-3 MΩ were filled with this solution and placed 10 μm away from the targeted myelin sheaths. These myelin sheaths were chosen to be at least 10 μm away from the soma to minimise the soma exposure to BAY 60-6583 application. The puff-pipette was placed approximately perpendicular to the targeted myelin sheaths (see Fig. 4e and 5a). Positive pressure was applied to the puff-pipette either with a micro-injector (PMI-100, Dagan) for 20 ms at 10 psi or controlled manually with a syringe. In both cases, positive pressure was calibrated and live monitored to eject drug over a radius of ≤20 μm (from the tip of the pipette set at a saturating intensity for Alexa Fluor 594, to the edge of detectable fluorescence; see also Supplementary Video 3), forming a sphere with a volume of ≤33,510 µm^3^. The flow of the perfusion was set to wash out the puffed drug in the direction away from the oligodendrocyte, further minimising the contact of the puffed solution with the soma and the other myelin sheaths located on the opposite side of the puff-pipette.

### Patch-clamping, calcium uncaging, and imaging of astrocytes

Astrocytes were detected by their morphology and by labelling with the selective marker sulforhodamine 101 (SR 101). Before the experiments, the slices were incubated with 1 μM SR101 (Tocris Bioscience) added to the slicing solution for 20 minutes at 37 °C. Microelectrodes with resistances of 7-8 MΩ were filled with a K-methylsulfate intracellular solution containing (in mM): 100 KMeSO_4_, 50 KCl, 2 MgCl_2_, 4 MgATP, 0.3 Na_2_GTP, and 10 HEPES (pH adjusted to 7.2 with KOH). Fluo-4 (100 μM; ThermoFisher Scientific) was added to image calcium changes and 5 mM NP-EGTA (ThermoFisher Scientific) was added to allow uncaging of calcium in single astrocytes. Images were acquired at 2 Hz with a confocal argon laser and NP-EGTA photolysis was obtained by applying 2-photon excitation at 720 nm to the soma (10 iterations, 2.55 µs pixel dwell time). Laser intensity was set at 10 mW and, if needed, was increased gradually until a calcium concentration rise was evoked^5^ (see also Supplementary Video 4). Intensities just below those needed to evoke a detectable [Ca^2+^]_i_ rise were used as control experiments to check that 2-photon illumination alone was not responsible for the effects observed. Node-astrocyte process distance was measured by drawing a line perpendicular to a myelinated axon. The line started at the centre of a node and ended at the closest astrocyte process localised in the same x-y plane as the node.

### Immunohistochemistry

Brains from Caspr-GFP mice were perfusion-fixed in 4% paraformaldehyde (PFA) in 0.01 M phosphate buffer saline (PBS) and fixed tissue was then cut into 70 μm-thick slices. Alternatively, 250 μm-thick acute slices from mouse were immersion-fixed in a solution containing 4% PFA, 4% sucrose, and 0.1M PBS. The slices were permeabilised with 0.2% Triton X-100 (Sigma-Aldrich) in a blocking solution (10% goat serum in 0.01 M PBS) for 1 h. The slices were then incubated overnight at 4 °C with the following primary antibodies, as required: rabbit anti-A_2b_R (Cohesion Bioscences, CPA3755, 1:100), chicken anti-GFP (Millipore, AB16901, 1:1000), mouse anti-NKCC1 (T4, Developmental Studies Hybridoma Bank, 1:100), rabbit anti-phospho-NKCC1 Thr212/Thr217 (Millipore, ABS1004, 1:200), rat anti-MBP (Millipore, MAB386, 1:200) and rabbit anti-GFAP (Millipore, AB5804, 1:500). The slices were then incubated for 2 hours at room temperature with the following secondary antibodies, as required (ThermoFisher Scientific, 1:500): anti-chicken Alexa Fluor 488, anti-rabbit or anti-mouse Alexa Fluor 546, and anti-rat Alexa Fluor 647. The slices were mounted in Dako Fluorescent Mounting Medium. Slices were washed with 0.01M PBS three times for 10 minutes between each step. Negative control experiments were also performed to check for any labelling caused by unspecific binding of the secondary antibodies: for that purpose, we followed the same protocol but omitting the incubation with the primary antibodies. Slices were imaged using a Zeiss LSM700 confocal microscope with a Zeiss Plan-Apochromat 63x oil immersion lens, and images were acquired with ZEN Microscope Software (Zeiss). Z-stacks of 1 μm-interval were imaged and their maximum intensity was projected using ImageJ (FIJI). To quantify the presence of A_2b_R, NKCC1 and pNKCC1 in the nodal regions, a line at least 10 µm-long centred on the node was drawn along the axons to plot intensity profiles, covering the whole length of the node and the paranodes, and 1-2 µm of the juxtaparanodes. The width of the line was adjusted to fit the width of the Caspr-labelled paranodes. We defined the proteins to be localised in the nodal regions if: (i) they were in contact with or overlapped the Caspr labelling and (ii) their mean intensities at these sites were >2, >3.5, or >5 times higher than A_2b_R, NKCC1 and pNKCC1 background intensity away from the axon, respectively (a higher threshold was needed for antibodies of lower quality).

### Live-detection of myelin sheaths with Fluoromyelin

Before the experiments, the slices were incubated with FluoroMyelin Red (ThermoFisher Scientific, 1:50) added to the slicing solution for 30 minutes at 37 °C. FluoroMyelin Red is a marker for compact myelin that was previously used on cell culture^10^ and *in vivo*^52^. Here, we used FluoroMyelin to image myelin in the corpus callosum of live brain slices in P14 mice. At this age, single myelinated axons can be traced because the density of myelinated axons is much lower than at later ages. To assess the association between Caspr and Fluoromyelin, we plotted intensity profiles of nodal regions, as detailed above, covering the entire length of the node and the Caspr-labelled paranodes and extended the profiles towards the juxtaparanodes (flanking the paranodes, away from the nodes of Ranvier). The width of the line was adjusted to fit the width of the Fluoromyelin labelling. Nodes of Ranvier were defined as myelinated if Fluoromyelin contacted Caspr. In some instances, Fluoromyelin could be seen away from a non-labelled paranode along the same axon, indicating that either myelination near this paranode was not completed or that myelin near this paranode was not compact enough to be labelled by Fluoromyelin.

### Estimating myelin sheath conductance

To estimate the myelinated process conductance per membrane area and the specific conductivity of internodal membrane from patch-clamping oligodendrocytes at the soma, several assumptions were made. We ignored the possible voltage non-uniformity between the soma and the inner and outer tongues of the myelin, as well as the contribution of the short processes connecting myelinated processes to the soma (both parameters are expected to have only a small effect on internode conductive properties^29^). We also assumed that all the membranes of the myelin have the same electrical properties, but only the outermost membrane is conductive because the extracellular space around all the inner layers is too restricted to allow significant current to leave the cell. The slope of the linear regression of the graph plotting the input conductance as a function of total internode length per cell was significantly different from zero (Extended Data Fig. 11a), indicating that internodes are electrically coupled to the soma as previously shown^29^. The slope of the linear regression gave the conductance per unit length of process in the absence of A_2b_R activation as 44.88 pS/µm. We then plotted the increase in input conductance for each cell obtained by puffing BAY 60-6583 onto myelin sheaths, as a function of the estimated sum of the lengths of the internodal processes targeted by the puff (Extended Data Fig. 11b). The drug was puffed over a radius of ≤20 μm as described above, and the lengths of the internodal processes contained within the sphere formed by the puff were summed. The linear regression for this graph was forced to go through the origin of the coordinates (as no puff or puffing aCSF should not have any effect; Extended Data Fig. 7f). The slope, which was also significantly different from zero, gave the A_2b_R-mediated rise in input conductance per unit length of process as 108.4 pS/µm. On average, uncaging-evoked astrocyte Ca^2+^ activity raised the conductance by 5.28 nS. If we assume that the puffed agonist and the adenosine generated by elevating astrocyte [Ca^2+^] activate A_2b_Rs to the same extent, then this (simplistically) corresponds to an effect on 48.71 µm of internode length per oligodendrocyte (5.28 nS divided by 108.4 pS/µm). Based on our assumption that current solely flows to the extracellular space across the outer membrane of myelin sheaths, the membrane area per unit length of an internode that provides the conductance is equal to 2π×(outer radius of myelin). In the computer model that we used (where the parameters were based on experimental data, see below), we assumed that the g-ratio is 0.81 and the internodal axon diameter is 0.73 µm, thus giving an outer diameter of 0.90 µm and a membrane area per 1 μm length of myelin of 2.83 μm^2^. An oligodendrocyte membrane area providing 44.88 pS (as defined by the slope of the linear regression in Extended Data Fig. 11a) would thus give a conductance density of 15.86 pS/μm^2^, which we assumed to be the baseline conductance density of each of the 10 membranes making up the 5 myelin layers in our computer model (see below). These estimates are similar to values used in previous computer simulations^1,29,53^, and are not expected to alter significantly the conduction speed compared to values assumed in earlier studies, as shown in Extended Data Fig. 11c and other works^29,53^. The A_2b_R-mediated rise in conductance density was calculated assuming that, for each 1 µm of myelin, the conductance increases by 108.4 pS to reach 153.28 pS (108.4 pS added to 44.88 pS), i.e. a 3.42-fold increase in the outer myelin membrane conductance per unit length of process. The conductance density of the outer layer would thus increase from 15.86 pS/μm^2^ to 54.16 pS/μm^2^ (153.28 pS divided by 2.83 μm^2^). For 10 membranes forming 5 myelin layers (with only the outer membrane increasing its conductance in response to adenosine), in order to correctly reproduce the overall change in trans-myelin sheath resistance when adenosine is added, the conductance density which is implemented in the computer model in all of the myelin membranes would have to increase by a factor of 1.07, from 15.86 pS/μm^2^ to 16.97 pS/μm^2^

### Computer simulation of infinite axon

As in our previous studies^1,5,29^, action potential conduction along myelinated axons was simulated using MATLAB. Electrophysiological parameters were based on the finite impedance double cable model (model C) of Richardson et al.^53^, except that the membrane capacitance^54^ was taken as the physiologically measured value of 0.9 μF/cm^2^. The differential equations of the model were derived and solved as in the myelinated axon model of Halter and Clark^55^, in which the axon is divided into compartments representing the node, paranode and internode. Conduction speed simulations of long callosal-like axons were carried out as previously^1^, where the parameters used were based on experimental data. The speed was measured between nodes 20 and 30 in a uniform axon containing 51 nodes and 50 internodes of constant lengths (nodes: 1.5 µm, internodes: 81.7 µm) and diameters (nodes: 0.64 µm, internodes: 0.73 µm). The periaxonal space thickness, which has an important effect on conduction speed^1,15^, was set at 15 nm, except in the paranodes (the 1.9 µm end parts of the internodes) where the width was reduced to 0.0123 nm^1^. Assuming a myelin wrap periodicity of 15.6 nm, 5 myelin wraps were needed to set the g-ratio close to 0.8. Each node expressed fast Na_V_, persistent Na_V_ and slow K_V_ channels at fully-activated conductance densities of 10, 0.01 and 0.4 mS/mm^2^ respectively. The node resting potential was set to −82 mV by adjusting the magnitude of a leak conductance (to 0.113 mS/mm^2^) with a reversal potential of −84 mV (the reversal potential of the other K^+^ currents in the model). The leak current represents K^+^ leak channels such as TRAAK present in the nodes of Ranvier^56,57^. To investigate the effect of A_2b_Rs/NKCC1 on conduction speed, we varied the node length and the myelin membrane conductance within a range of values that includes our experimental observations^1,29^: from 0.01 µm to 10 µm and from 1 pS/μm^2^ to 150 pS/μm^2^, respectively. When varying node length, internodal lengths were shortened accordingly and the nodal channel number was kept constant to represent the A_2b_R/NKCC1-mediated adaptations observed, or the nodal channel density was kept constant to simulate channels being inserted into or removed from the nodal membrane. When channel density was kept constant, node lengths below 0.5 µm did not generate action potentials. To test the conduction delay triggered by the effect of adenosine on one node, we measured the delay in action potential arrival at node 40 when increasing the length of node 20 from the shortest to the longest value observed (to mimic A_2b_R activation) and adding an adenosine-mediated I_h_ current (conductance changed from 0 to 0.1565 mS/mm^2^ to mimic A_2a_R activation^5^).

### Statistics

Statistical analyses and graphs were performed with GraphPad Prism 8. Data are presented as mean±SEM. To compare the percentages of motile versus stable nodes, elongating versus shortening nodes, and NKCC1/pNKCC1 detection in nodal regions, statistical comparisons were performed with the Chi-square goodness of fit test (to compare observed versus expected outcomes, for example testing the effect of a drug versus aCSF) or Chi-squared test (to assess whether categorical variables differ significantly, for example comparing two different age groups). When more than two groups were compared, P values were corrected for multiple comparisons using a procedure equivalent to the Holm-Bonferroni method (for N comparisons, the most significant P value is multiplied by N, the 2^nd^ most significant by N – 1, the 3^rd^ most significant by N – 2, etc.). For the other data, which are continuous variables, data normality was assessed with D’Agostino & Pearson omnibus or Kolmogorov-Smirnov tests. Comparisons of two normally distributed groups were made using two-tailed Student *t*-tests (paired or unpaired, where appropriate). When treatments involved more than two independent groups, statistical comparisons were performed with one-way or two-way ANOVA, and *post hoc* Tukey’s or Benjamini, Krieger, and Yekutieli’s multiple comparison corrections. Data that were not normally distributed were analysed with Wilcoxon matched-pairs signed rank tests. We assessed the difference between two slopes of linear regressions by using the *t*-statistic for the slope. P values are significant if they are less than 0.05, statistical units *n* are nodes unless mentioned. This was appropriate because for major results the variance within individual mice was greater than the variance between mice, for example in the expression of A_2b_Rs (10.6 times higher) and NKCC1 (9.8 times higher) in nodal regions.

## Data availability

All data are available in the manuscript or the supplementary materials. Code used for simulations is freely available from https://github.com/AttwellLab/MyelinatedAxonModel.

## Acknowledgements

This work was supported by European Research Council (BrainEnergy) and Wellcome Trust Awards (099222/Z/12/Z and 219366/Z/19/Z), and funding from the BHF/UK-DRI Centre for Vascular Dementia Research to D.A., an MRC PhD studentship to C.T., and an EMBO fellowship (ALTF 430-2019) to J.L.

## Author contributions

We thank B. Clark, D. Kullmann and G. Schiavo for comments on the manuscript. J.L. performed the electrophysiological experiments and computer simulations. J.L., T.Q., N.W, S.M. and C.T performed the live-imaging and immunohistochemistry experiments. J.L. conceived the project with the guidance of D.A. J.L wrote the first draft of the manuscript and all other authors commented on the manuscript.

## Competing interests

The authors declare no competing financial interests.

## Extended data

**Extended Data Fig. 1:**
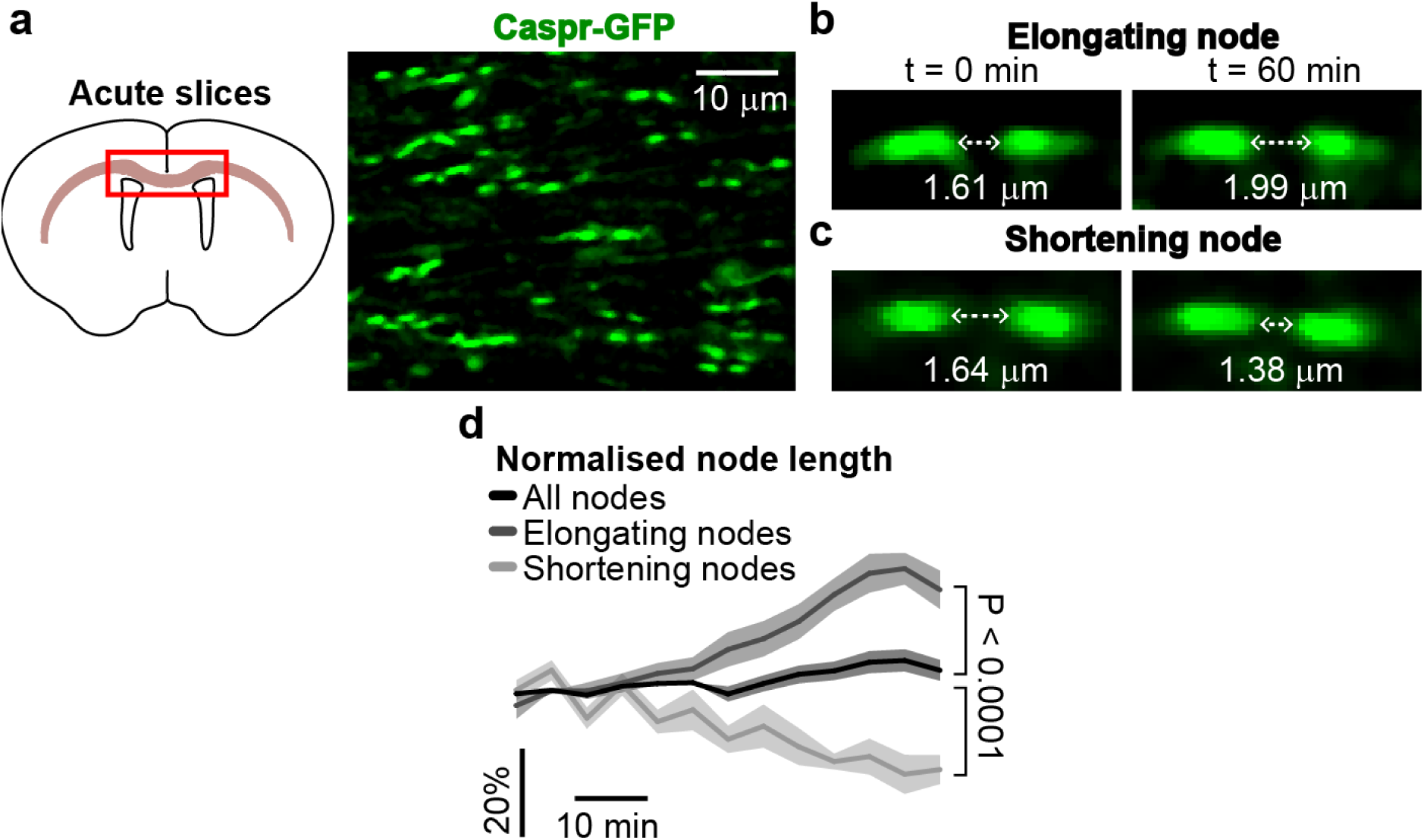
Node of Ranvier motility in live acute brain slices. **a** Nodes in the white matter projections of the corpus callosum imaged in live acute brain slices. **b-c** As in intact brains of anesthetised mice, some nodes could be seen elongating (**b**) and others shortening (**c**) in the white matter. **d** Percentage of change in node length, imaging nodes in slices every 5 minutes for one hour, for the elongating and shortening nodes, and for all the nodes together (*n*=17, 10 and 60, respectively). Average node length values of last 15 minutes are significantly different (P<0.0001, one-way ANOVA).

**Extended Data Fig. 2:**
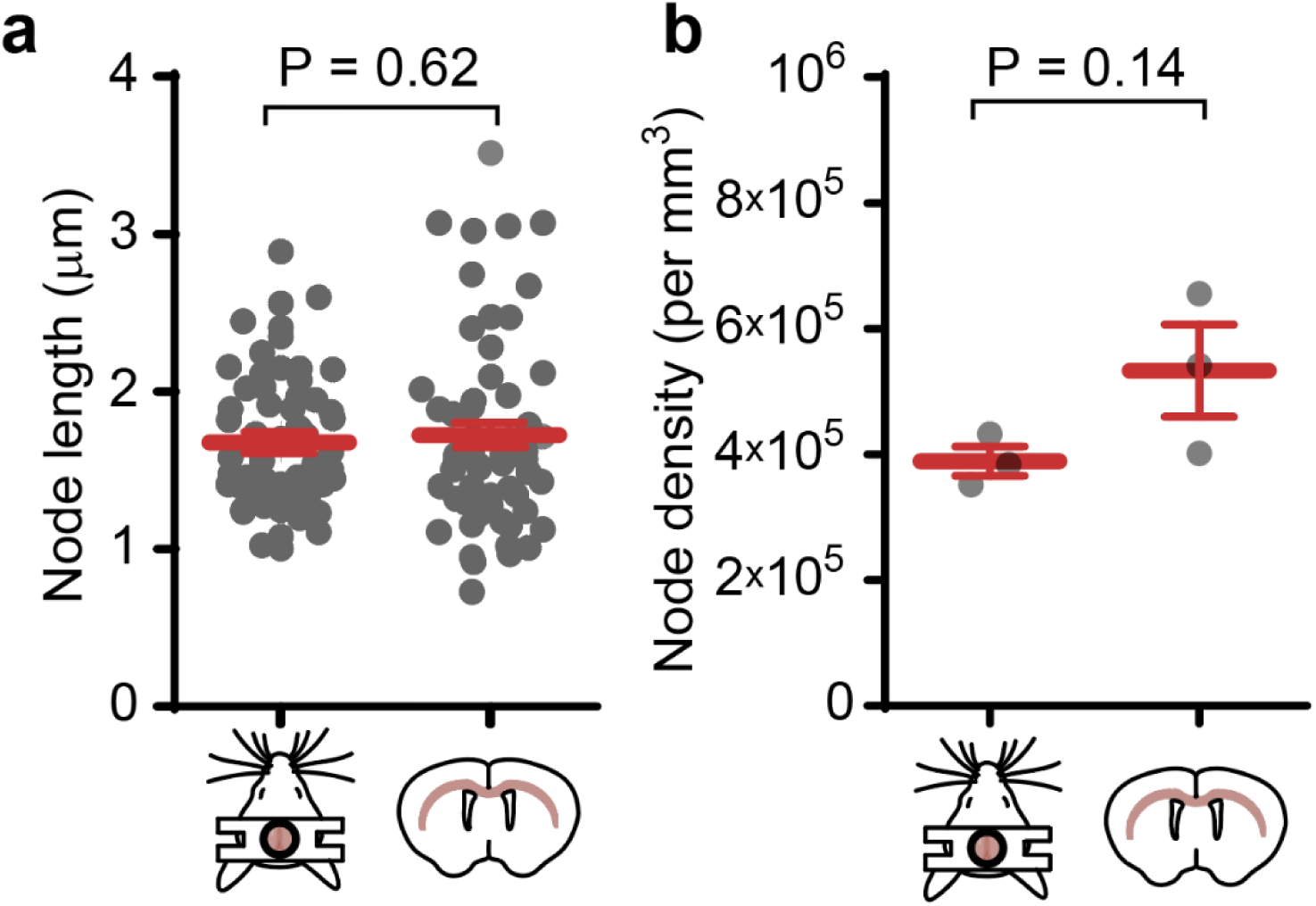
Nodes of Ranvier length and density are similar *in vivo* and *ex vivo*. a,b. Total mean node length (**a**; *n*=63 nodes *in vivo* and 60 *ex vivo*, unpaired t-test) and node density (**b**; *n*=3 mice *in vivo* and *ex vivo*, unpaired t-test) were similar when imaged through cranial window preparations in intact brains of live anesthetised mice (left) and in live coronal acute brain slices (right). Although not statistically significant, the apparent difference in node density (higher by 143,668 nodes/mm^3^ in slices) may arise from differences between the two types of white matter projections imaged: the cingulum *in vivo* and the corpus callosum *ex vivo*.

**Extended Data Fig. 3:**
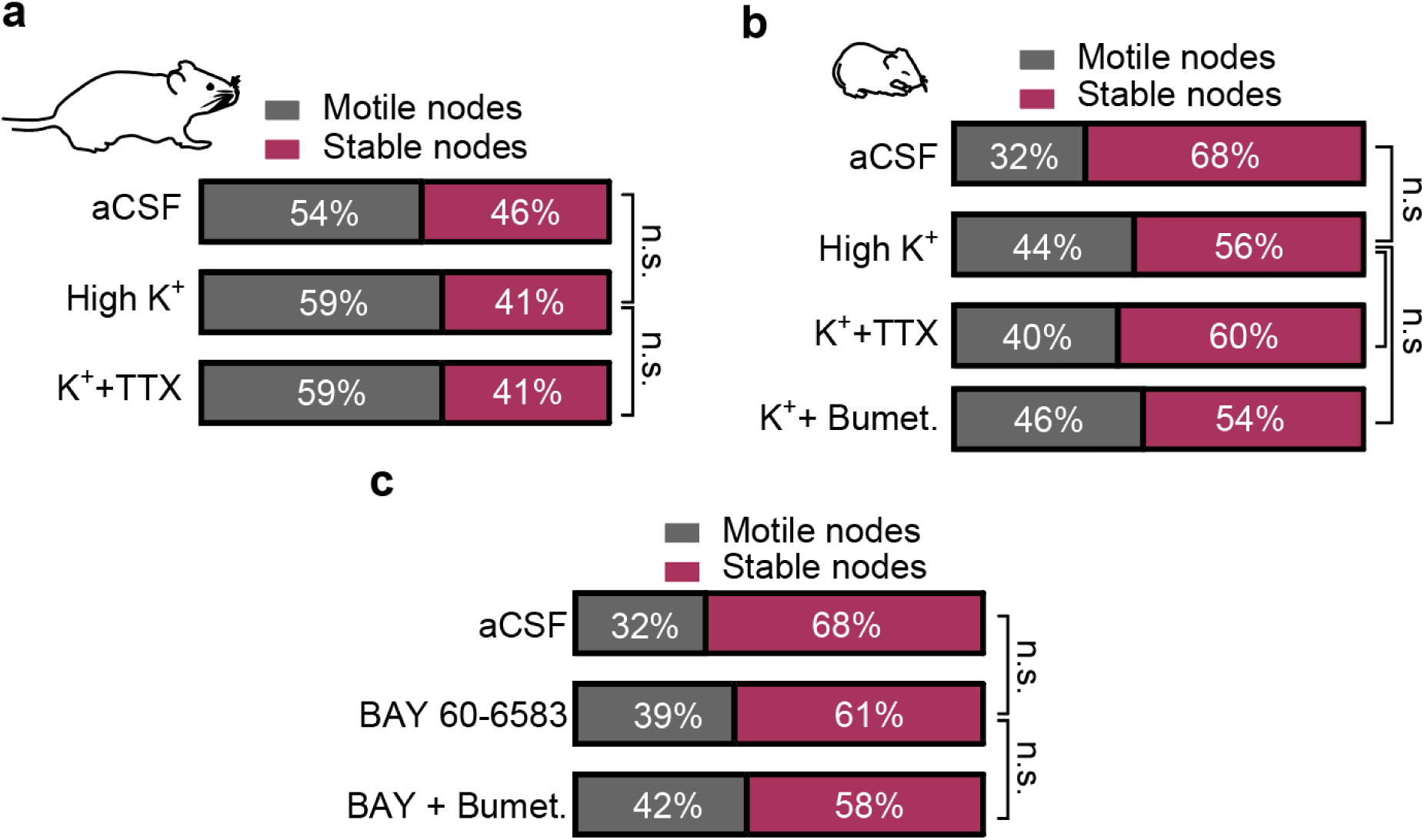
Fraction of motile nodes remains unchanged in different conditions. **a** In adult mice, high [K^+^]_o_, either with or without TTX, did not change the fraction of motile nodes (aCSF versus K^+^: P=0.32; High K^+^ versus K^+^ + TTX: P=1). **b** In young mice, high [K^+^]_o_, either with or without TTX and with or without bumetanide, did not change the fraction of motile nodes (aCSF versus High K^+^: P=0.051; High K^+^ versus K^+^ + TTX: P=0.84; K^+^ versus K^+^ + Bumetanide: P=0.69). **c** In young mice, superfusion of BAY 60-6583, either with or without bumetanide, did not change the fraction of motile nodes (aCSF versus BAY 60-6583: P=0.27; BAY 60-6583 versus BAY 60-6583 + Bumetanide: P=0.54). n.s.: non-significant.

**Extended Data Fig. 4:**
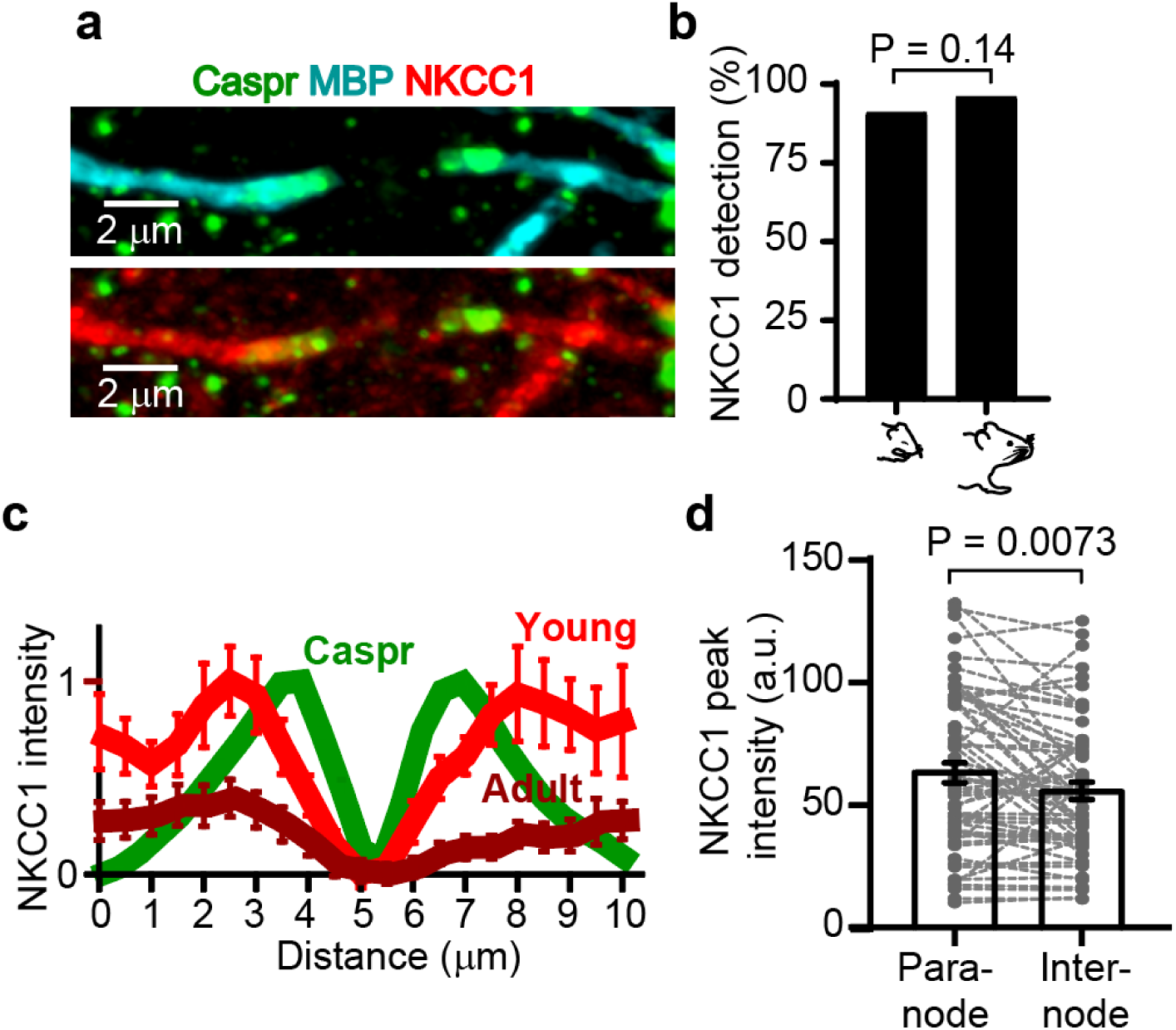
NKCC1 fluorescence detection along myelinated axons. **a** NKCC1 (red) colocalises with the paranodal marker Caspr (green) and the myelin marker MBP (cyan), and flanks the nodes of Ranvier on both sides. **b** NKCC1 was detected in the paranodal regions of young and adult mice (*n=*75 and 103, respectively; P=0.14). **c** Mean of 20 NKCC1 intensity profiles from young mice (light red) and 20 NKCC1 intensity profiles from adult mice (dark red) normalised to the maximum mean peak intensity in young mice. Mean of 74 Caspr intensity profiles is shown to assess the positioning of NKCC1 around the nodes. **d** NKCC1 peak intensity in 62 young paranodes and internodes (paranode: 64.11±4.13 a.u. versus internode: 56.24±3.51 a.u; P=0.0073, paired *t*-test).

**Extended Data Fig. 5:**
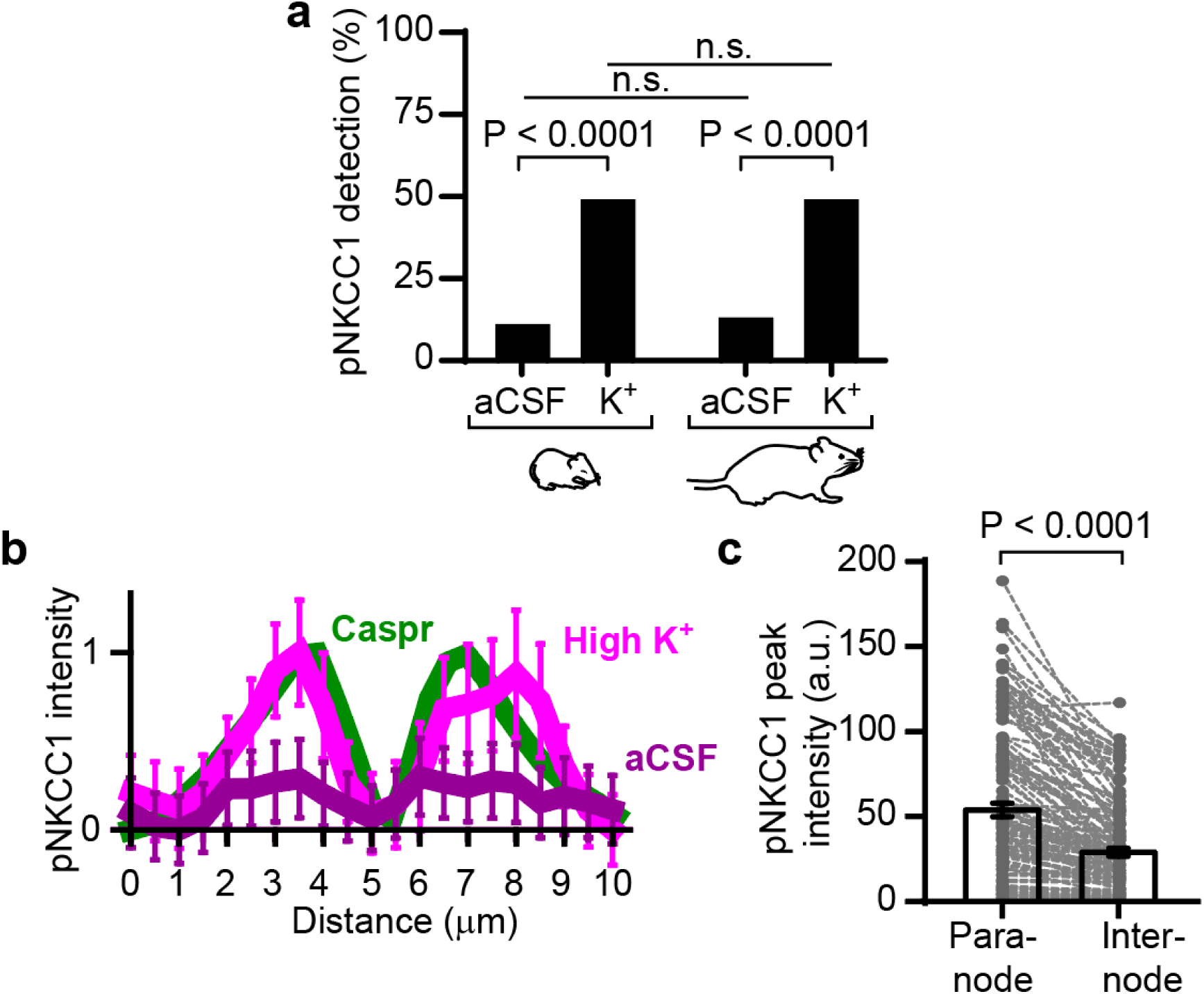
pNKCC1 fluorescence detection in paranodes. **a** Exposing slices to high K^+^ raised the presence of the phosphorylated form of NKCC1 (pNKCC1; a detectable pNKCC1 signal was defined as being at least 5-fold larger than background) in the paranodal regions of both young and adult mice (young mice: *n*=120 in aCSF and 127 high K^+^, P<0.0001; adult mice: *n*=90 in aCSF and 133 high K^+^, P<0.0001). The fraction of pNKCC1 present in paranodal regions was similar in young and adult slices exposed to the same conditions (n.s.: non-significant). **b** Mean of 20 pNKCC1 intensity profiles from slices incubated in high [K^+^]_o_ (light magenta) and 20 pNKCC1 intensity profiles from slices incubated in aCSF (dark magenta) normalised to the maximum mean peak intensity in high [K^+^]_o_-incubated slices. pNKCC1 peak fluorescence is concentrated in the paranodal regions, colocalising with Caspr and flanking the nodes. **c** pNKCC1 peak intensity in 126 young paranodes and internodes incubated in high [K^+^]_o_ (paranode: 54.74±4.02 a.u. versus internode: 29.87±2.36 a.u; P<0.0001, paired *t*-test).

**Extended Data Fig. 6:**
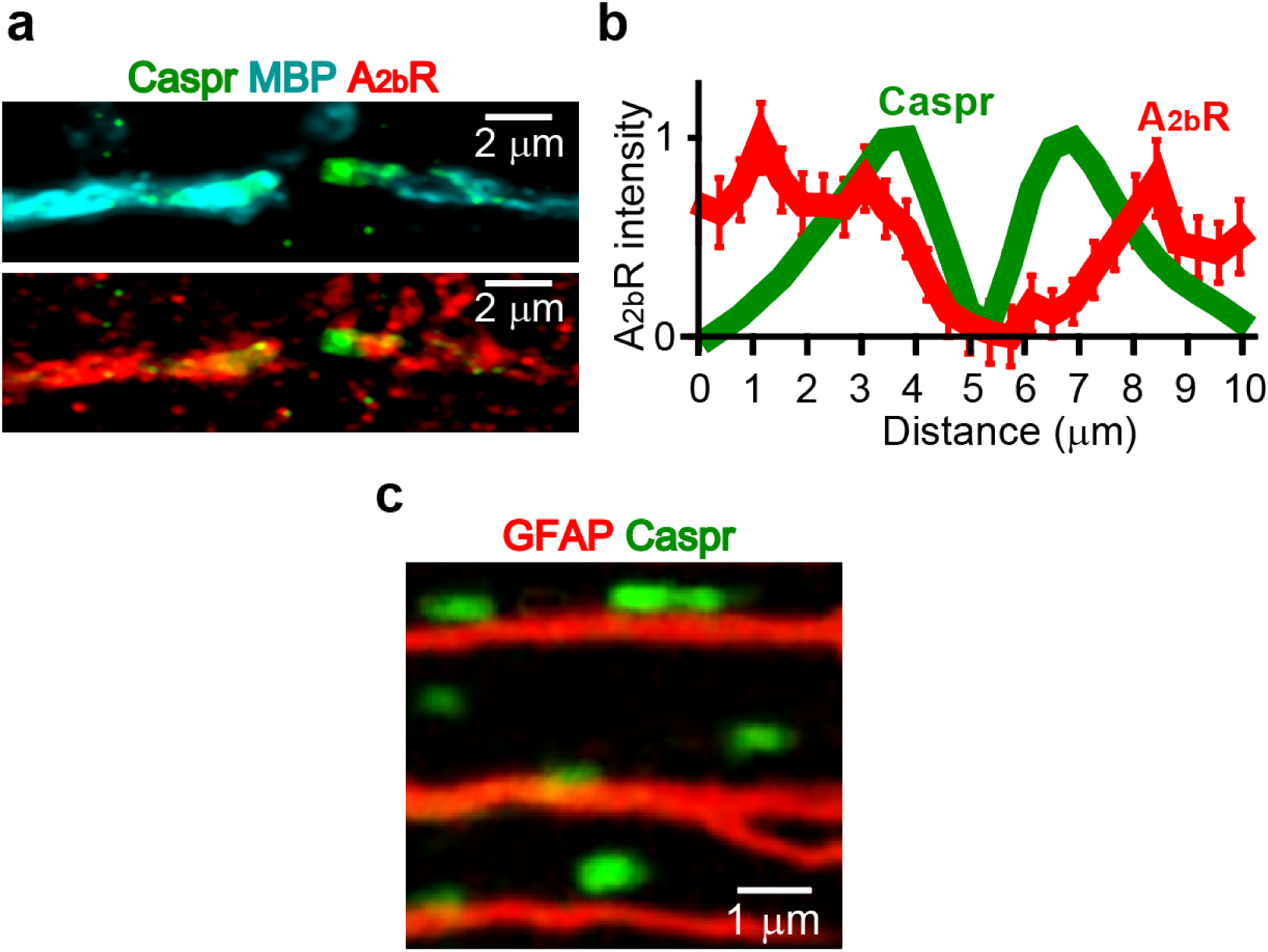
A_2b_R expression along myelinated axons and astrocyte process association with nodes. **a** A_2b_Rs (red) colocalise with the paranodal marker Caspr (green) and the myelin marker MBP (cyan), and flank the nodes of Ranvier on both sides. **b** Mean of 50 A_2b_R (red) and 74 Caspr (green) intensity profiles showing the presence of A_2b_Rs in the paranodal and internodal sections of the myelinated axons, flanking the nodes of Ranvier. **c** GFAP-labelled astrocyte processes (red) run parallel to myelinated axons, and thus are in close proximity to nodes, paranodes and internodes, as shown by the association with Caspr-labelled paranodes flanking three callosal nodes (green).

**Extended Data Fig. 7:**
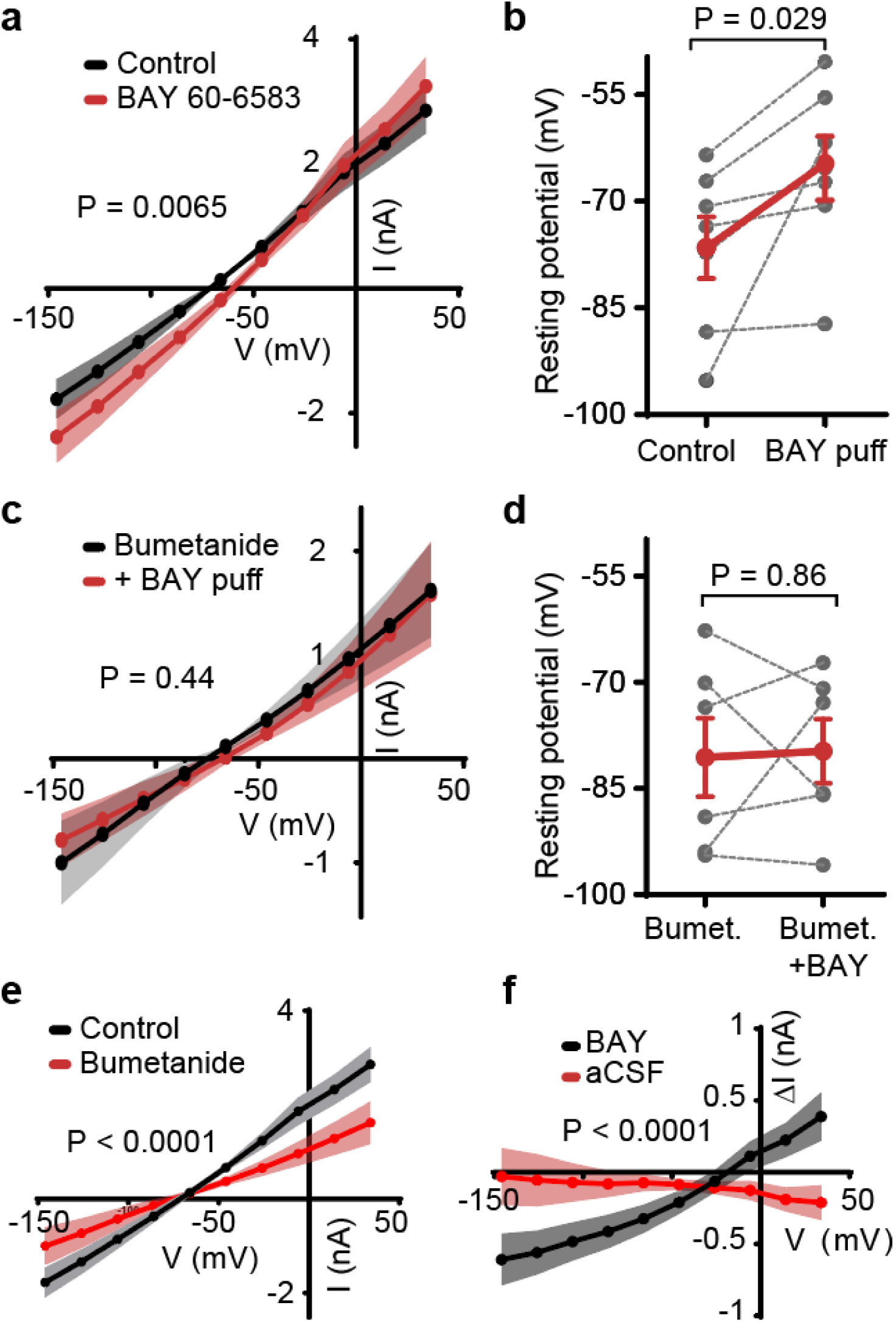
A_2b_R and NKCC1 on myelin sheaths depolarise the resting potential of oligodendrocytes. **a** I-V relation of 7 oligodendrocytes before and after puff-application of the A_2b_R agonist BAY 60-6583 onto dye-filled internodes (slopes are significantly different: P=0.0065). **b** Depolarising shift in the membrane resting potential following BAY 60-6583 puff (*n*=7; two-way ANOVA over panels b and d). **c** As in (**a**), but while including the NKCC1 blocker bumetanide within the aCSF. I-V relation of 6 cells did not change when puffing BAY 60-6583. **d** BAY puff did not evoke a depolarising shift in resting potential of oligodendrocytes when bumetanide was added to the aCSF (*n*=6; two-way ANOVA). **e** I-V relations of 7 oligodendrocytes in control conditions and 6 oligodendrocytes when bumetanide was bath-applied. **f** In contrast to BAY 60-6583 application, puffing aCSF does not vary membrane currents (red curve; P<0.0001 when comparing the slopes; slope of red curve is not significantly different from zero: P=0.14).

**Extended Data Fig. 8:**
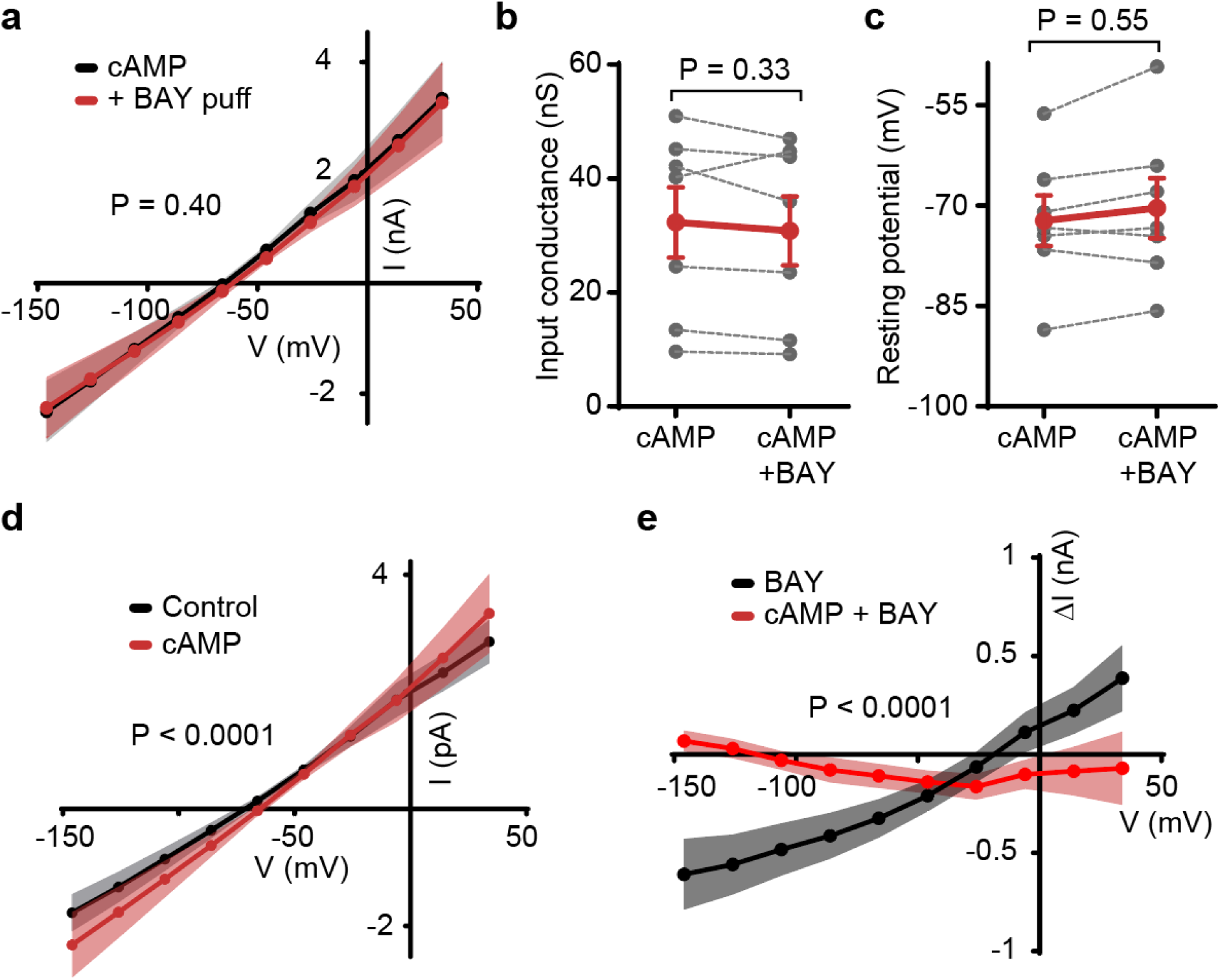
Intracellular cAMP mediates the effect of myelin A_2b_Rs on membrane current. **a** I-V relation of 7 oligodendrocytes loaded with cAMP before and upon puff-application of BAY 60-6583 on myelin sheaths. Black and red slopes are similar (P=0.40). **b-c** Input conductance (**b**) and resting potential (**c**) of oligodendrocytes loaded with cAMP before and after BAY 60-6583 application (P=0.33 and 0.55, two-way ANOVA with Fig. 4g and Extended Data Fig. 7b, respectively; *n*=7). **d** I-V relations of 7 oligodendrocytes in control conditions and 7 oligodendrocytes when cAMP is loaded intracellularly. **e** Puff-application of the A_2b_R agonist BAY 60-6583 did not vary the membrane currents when oligodendrocytes and their myelin sheaths included 50 µM cAMP (P<0.0001 when comparing the slopes).

**Extended Data Fig. 9:**
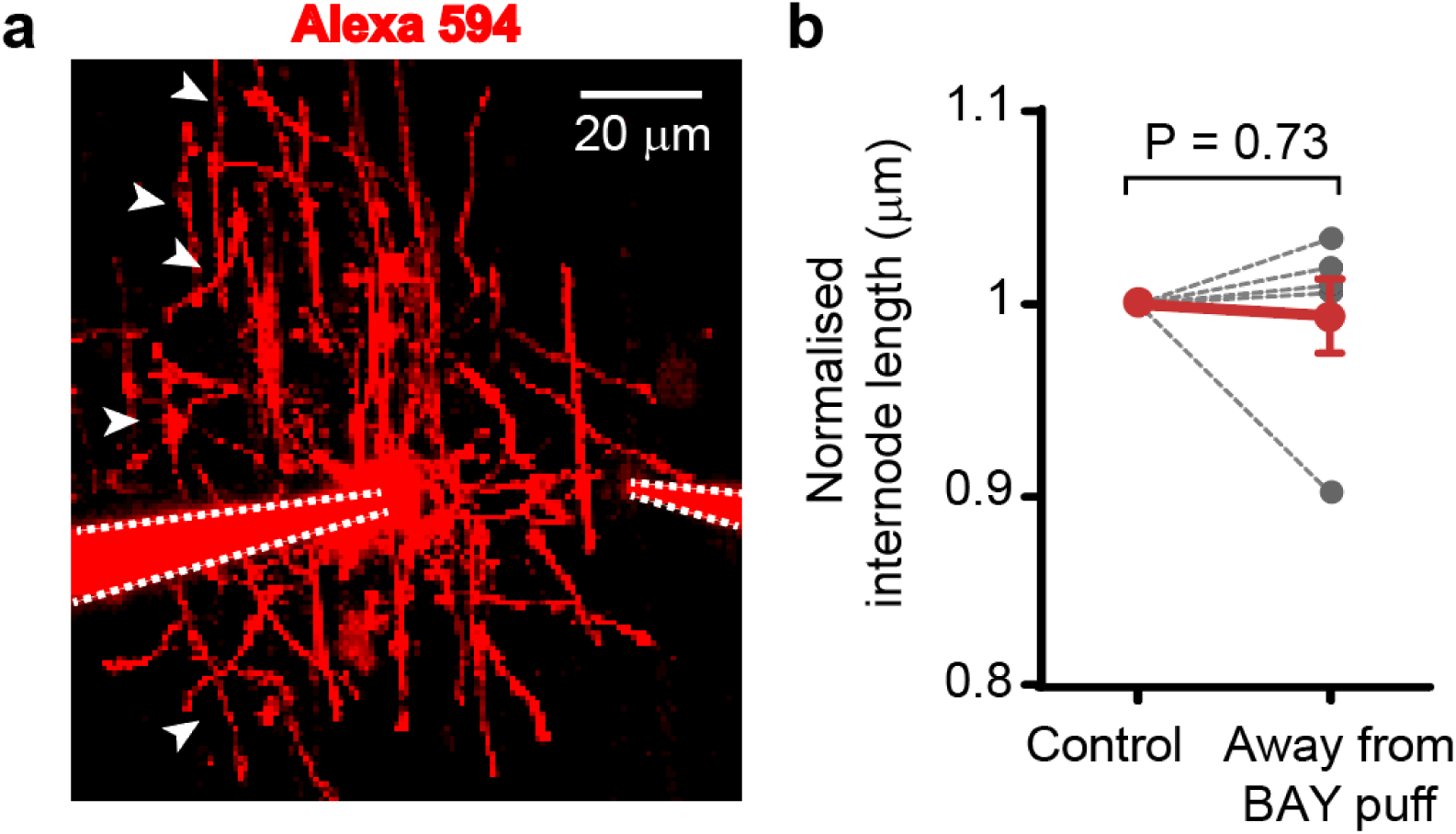
Mean internode length is stable away from BAY 60-6583 application. **a** Specimen oligodendrocyte patched-clamped and dye-filled with Alexa 594 in the left pipette. White arrowheads show internodes emerging from the oligodendrocyte that were not targeted by BAY 60-6583 local application (see Methods and Supplementary Video 3). **b** Normalised changes in internode length observed on the side opposite to and upstream of the puff-pipette, away from the area targeted by BAY 60-6583 local application (see Methods; n=6, two-way ANOVA).

**Extended Data Fig. 10:**
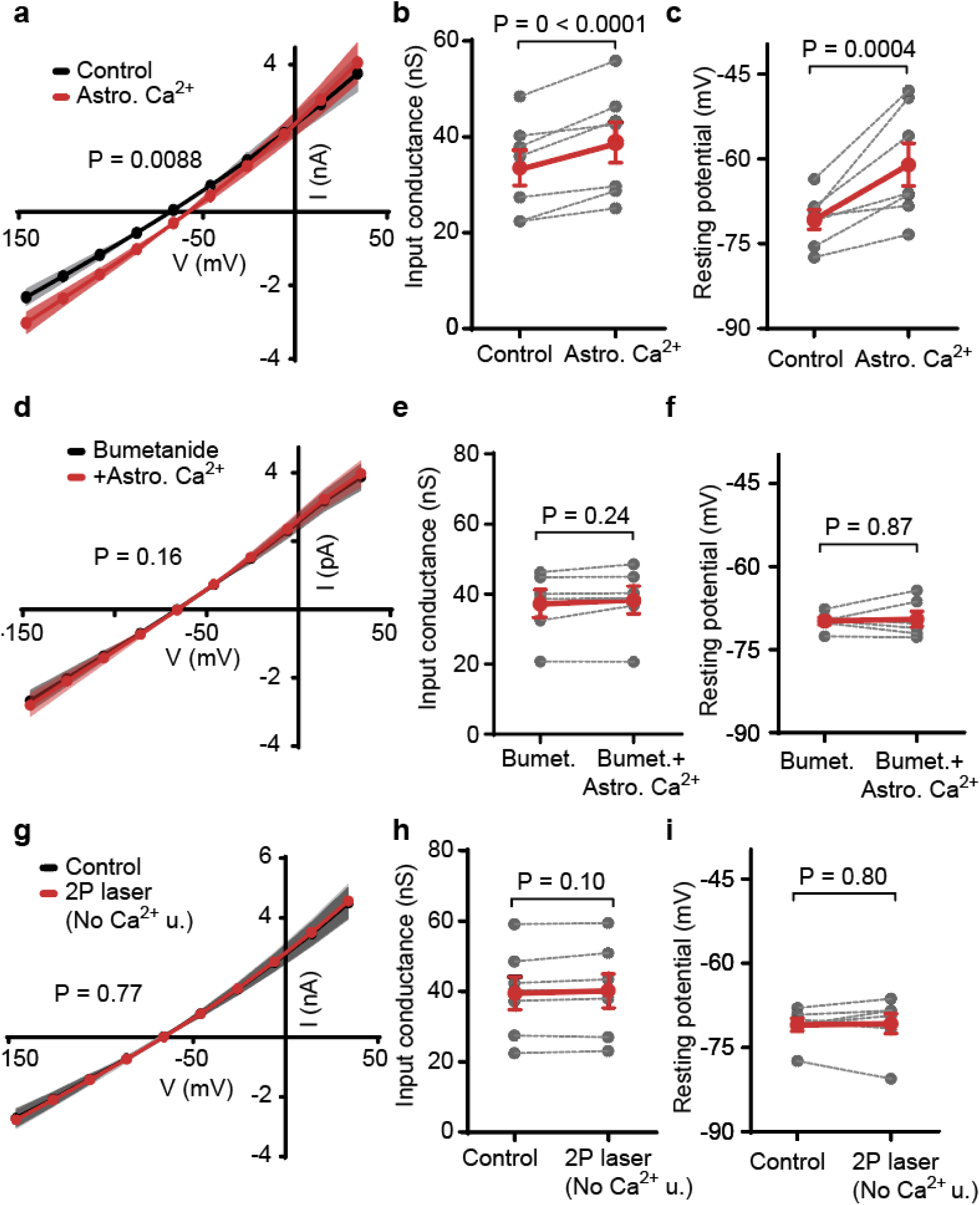
Astrocyte Ca^2+^ rises evoke NKCC1-mediated variations in oligodendrocyte membrane current. **a** I-V relation of 7 oligodendrocytes before and after uncaging astrocyte Ca^2+^ near a patch-clamped oligodendrocyte (slopes are significantly different: P=0.0088). **b-c** Input conductance (**b**; two-way ANOVA over panels b and e, *g_i_*_n_ values were normalised to control for comparisons) and resting potential (**c**) of oligodendrocytes before and after uncaging astrocyte Ca^2+^ (*n*=7; two-way ANOVA over panels c and f). **d** I-V relation of 6 oligodendrocytes before and after uncaging astrocyte Ca^2+^ when bumetanide was included to the external solution (slopes are similar: P=0.16). **e-f** Input conductance (**e**) and resting potential (**f**) of oligodendrocytes before and after uncaging astrocyte Ca^2+^ in the presence of bumetanide in the aCSF (*n*=6; two-way ANOVA). **g** I-V relation of 7 oligodendrocytes before and after two-photon illumination that did not elicit a Ca^2+^ rise [2P laser (No Ca^2+^ u.)] (slopes are similar: P=0.77). **h-i** Input conductance (**h**) and resting potential (**i**) of oligodendrocytes before and after two-photon illumination that did not elicit a Ca^2+^ rise (*n*=7).

**Extended Data Fig. 11:**
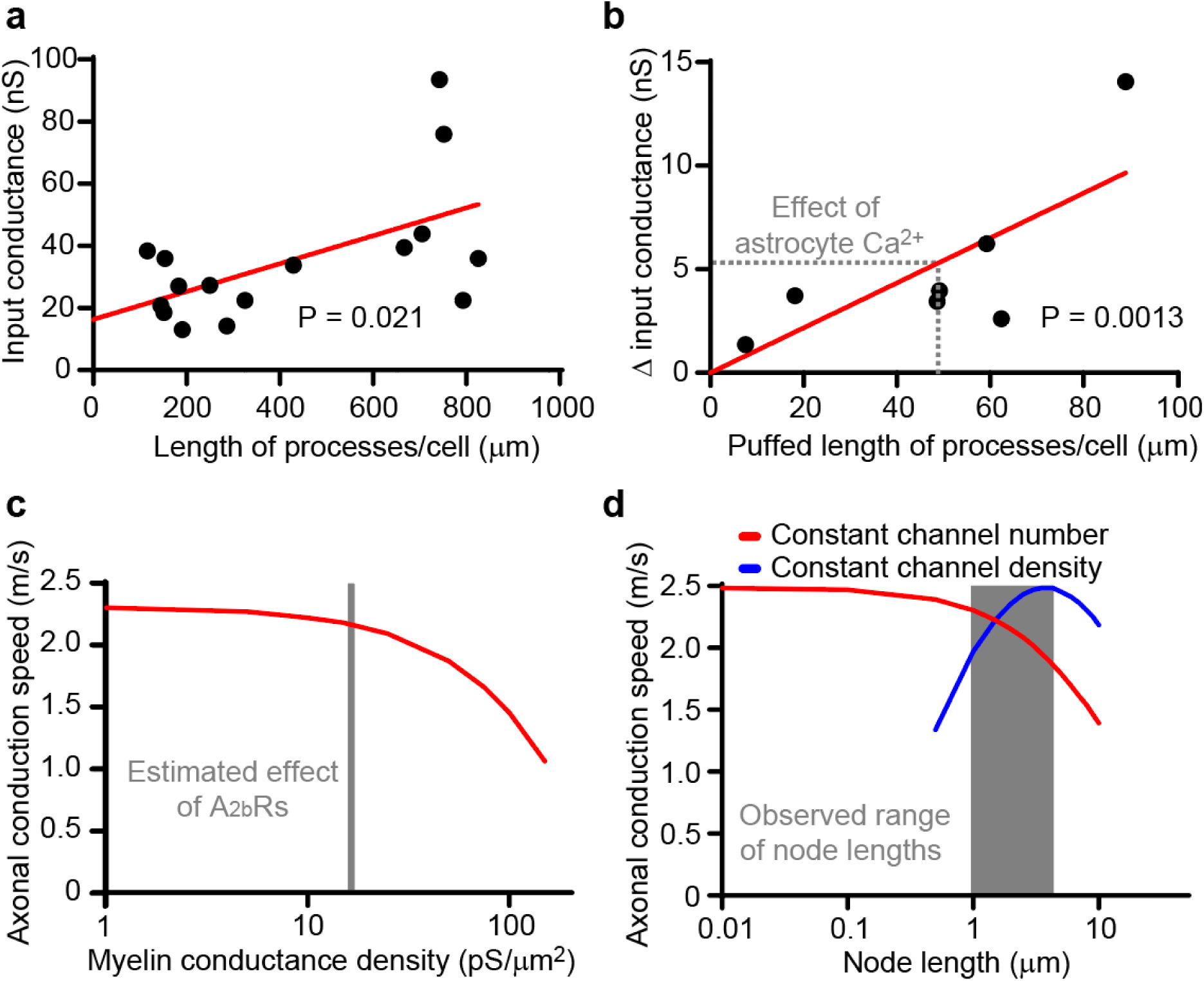
Predicted effects of adaptations in myelin conductance and node length on axonal conduction speed. **a** Input conductances of 16 patch-clamped oligodendrocytes as a function of the total length of their internode processes. Red line shows linear regression with R^2^=0.33; the slope (44.88 pS/µm), which gives the conductance per unit length of myelin sheath, is significantly different from zero (P=0.021). **b** Increase in input conductances of 7 patch-clamped oligodendrocytes as a function of the sum of all the internode process lengths onto which BAY 60-6583 was puffed (see Methods). Red line shows linear regression with R^2^=0.58; the slope (108.4 pS/µm), which gives the rise in myelin conductance density evoked by A_2b_R activation, is significantly different from zero (P=0.0013). Grey dashed lines show the mean rise in oligodendrocyte conductance obtained by uncaging Ca^2+^ in astrocytes (5.28 nS) and the estimated length of processes that would trigger this rise if targeted by BAY 60-6583 (48.71 µm) (see Discussion and Methods). **c** Predicted impact of myelin conductance density on axonal conduction speed in computer simulations. Grey area shows the estimated increase in myelin conductance density when myelin A_2b_Rs are activated in young mice, rising from 15.86 pS/μm^2^ to 16.97 pS/μm^2^ (see Methods). **d** Predicted impact of node length variations on axonal conduction speed if nodal channel number is kept constant, simulating a longitudinal shortening or lengthening of the paranodal ends of the internodes via A_2b_R activity (red curve), or if nodal channel density is kept constant, simulating the insertion or removal of membranes with channels at the nodes (blue curve, the speed starts to decrease from 4.4 µm). Grey area shows the range of node lengths observed in the white matter of 2-week-old mice: between 0.96 µm and 4.35 µm.

**Extended Data Fig. 12:**
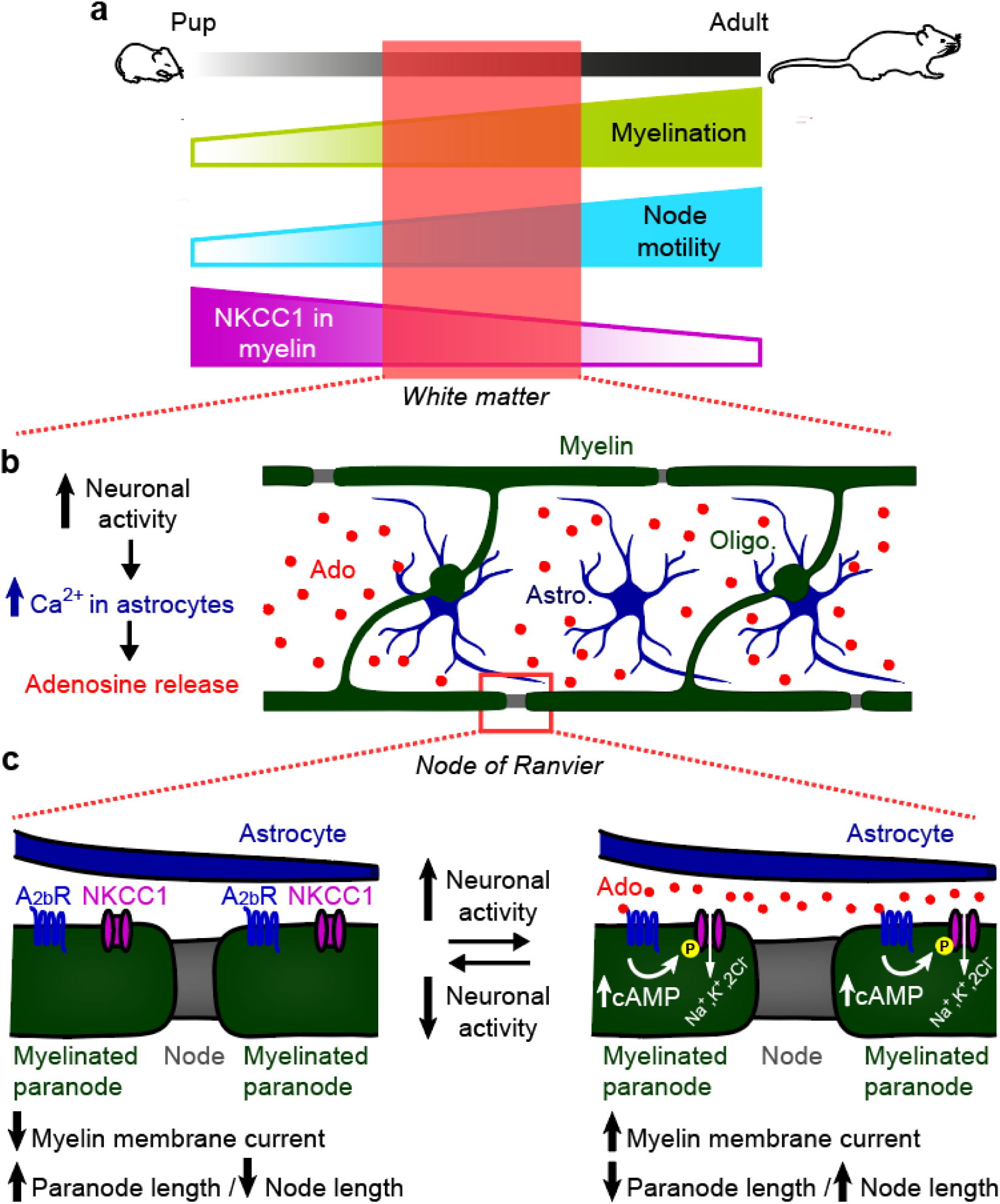
Mechanisms regulating node of Ranvier motility in the white matter. **a** Diagram summarising the changes observed in the white matter. The fraction of motile nodes increased as myelination progressed and node density rose, suggesting that myelination may confer motility to the nodes. In juvenile mice, NKCC1 co-transport expression is upregulated in myelin sheaths flanking nodes of Ranvier. **b** In the white matter, oligodendrocytes (Oligo.) form myelin sheaths and astrocyte (Astro.) processes associate with myelinated axons at nodes and internodes. Neuronal activity promotes astrocyte Ca^2+^ activity which in turn elicits ATP release, and ATP is converted to adenosine (Ado) in the extracellular space. We found in this study that node and paranode structural dynamics are linked to neuronal activity and astrocyte Ca^2+^ rises. **c** Diagram depicting the cellular mechanisms mediating node motility in the white matter. In response to high neuronal activity and astrocyte Ca^2+^ rises near myelinated axons, adenosine release activates A_2b_Rs on the myelin, which raises intracellular cAMP. This increases internodal membrane currents and phosphorylates NKCC1 at the myelinated paranodes, leading to a shortening of the paranodal ends of the internodes, thereby elongating nodes of Ranvier.

**Supplementary Video 1: Node of Ranvier elongating over time.**

Time-lapse images taken every five minutes for one hour of a node of Ranvier elongating in the corpus callosum of an adult mouse, as shown in Extended Data Fig. 1 and as defined in the Methods. White sign shows node length at timepoint t=0 superimposed on the time-lapse images to highlight the changes in length.

**Supplementary Video 2: Node of Ranvier shortening over time.**

Time-lapse images taken every five minutes for one hour of a node of Ranvier shortening in the corpus callosum of an adult mouse, as shown in Extended Data Fig. 1 and as defined in the Methods. White sign shows node length at timepoint t=0 superimposed on the time-lapse images to highlight the changes in length.

**Supplementary Video 3: Puff-application of BAY 60-6583 on myelin sheaths.**

An oligodendrocyte in the corpus callosum of a young mouse was patch-clamped and dye-filled with Alexa Fluor 594, which diffuses into the myelin sheaths emerging from the same oligodendrocyte. Another pipette was used to puff-apply the A_2b_R agonist BAY 60-6583 onto myelin sheaths. The puff-pipette included Alexa Fluor 594 to delineate the region targeted by the puff-application. Time-lapse images show the spread of the puff-application over time.

**Supplementary Video 4: Ca^2+^ uncaging in an astrocyte near myelin sheaths.**

Time-lapse images of an astrocyte in which Ca^2+^ was uncaged at the soma by two-photon excitation at 720 nm targeted at its soma. Ca^2+^ is seen propagating in the astrocyte processes, including those associating with myelin sheaths dye-filled with Alexa Fluor 594 (red). The astrocyte was loaded with NP-EGTA and Fluo-4 to uncage Ca^2+^ and monitor its rise, as shown in green. Time-lapse images of the astrocyte were superimposed on a still image of the patch-clamped oligodendrocyte dye-filled in red to highlight the position of the astrocyte processes and the myelin sheaths.

## Notes

### Competing Interest Statement

The authors have declared no competing interest.

### Summary of Updates

Computer simulations have been updated to reproduce the overall change in trans-myelin sheath conductance when myelin adenosine A2b receptors are activated (Extended Data Fig. 11c).

